# Cluster-specific gene markers enhance *Shigella* and Enteroinvasive *Escherichia coli in silico* serotyping

**DOI:** 10.1101/2021.01.30.428723

**Authors:** Xiaomei Zhang, Michael Payne, Thanh Nguyen, Sandeep Kaur, Ruiting Lan

**Affiliations:** School of Biotechnology and Biomolecular Sciences, University of New South Wales, Sydney, New South Wales, Australia

**Keywords:** Phylogenetic clusters, cluster-specific gene markers, *Shigella*/EIEC serotyping

## Abstract

*Shigella* and enteroinvasive *Escherichia coli* (EIEC) cause human bacillary dysentery with similar invasion mechanisms and share similar physiological, biochemical and genetic characteristics. The ability to differentiate *Shigella* and EIEC from each other is important for clinical diagnostic and epidemiologic investigations. The existing genetic signatures may not discriminate between *Shigella* and EIEC. However, phylogenetically, *Shigella* and EIEC strains are composed of multiple clusters and are different forms of *E. coli.* In this study, we identified 10 *Shigella* clusters, 7 EIEC clusters and 53 sporadic types of EIEC by examining over 17,000 publicly available *Shigella*/EIEC genomes. We compared *Shigella* and EIEC accessory genomes to identify the cluster-specific gene markers or marker sets for the 17 clusters and 53 sporadic types. The gene markers showed 99.63% accuracy and more than 97.02% specificity. In addition, we developed a freely available *in silico* serotyping pipeline named *Shigella* EIEC Cluster Enhanced Serotype Finder (ShigEiFinder) by incorporating the cluster-specific gene markers and established *Shigella*/EIEC serotype specific O antigen genes and modification genes into typing. ShigEiFinder can process either paired end Illumina sequencing reads or assembled genomes and almost perfectly differentiated *Shigella* from EIEC with 99.70% and 99.81% cluster assignment accuracy for the assembled genomes and mapped reads respectively. ShigEiFinder was able to serotype over 59 *Shigella* serotypes and 22 EIEC serotypes and provided a high specificity with 99.40% for assembled genomes and 99.38% for mapped reads for serotyping. The cluster markers and our new serotyping tool, ShigEiFinder (https://github.com/LanLab/ShigEiFinder), will be useful for epidemiologic and diagnostic investigations.

**Impact statement:** The differentiation of *Shigella* strains from enteroinvasive *E. coli* (EIEC) is important for clinical diagnosis and public health epidemiologic investigations. The similarities between *Shigella* and EIEC strains make this differentiation very difficult as both share common ancestries within *E. coli*. However, *Shigella* and EIEC are phylogenetically separated into multiple clusters, making high resolution separation using cluster specific genomic markers possible. In this study, we identified 17 *Shigella* or EIEC clusters including five that were newly identified through examination of over 17,000 publicly available *Shigella* and EIEC genomes. We further identified an individual or a set of cluster-specific gene markers for each cluster using comparative genomic analysis. These markers can then be used to classify isolates into clusters and were used to develop an *in silico* pipeline, ShigEiFinder (https://github.com/LanLab/ShigEiFinder) for accurate differentiation, cluster typing and serotyping of *Shigella* and EIEC from Illumina sequencing reads or assembled genomes. This study will have broad application from understanding the evolution of *Shigella*/EIEC to diagnosis and epidemiology.

**Data summary:** Sequencing data have been deposited at the National Center for Biotechnology Information under BioProject number PRJNA692536.

**Repositories:** Raw sequence data are available from NCBI under the BioProject number PRJNA692536.

## Introduction

*Shigella* is one of the most common etiologic agents of foodborne infections worldwide and can cause diarrhea with a very low infectious dose (1, 2). The infections can vary from mild diarrhea to severe bloody diarrhea referred to as bacillary dysentery. The estimated cases of *Shigella* infections are 190 million with at least 210,000 deaths annually, predominantly in children younger than 5 years old in developing countries (3–7). *Shigella* infections also have a significantly impact on public health in developed countries although most cases are travel-associated (8).

The *Shigella* genus consists of four species, *Shigella sonnei, Shigella flexneri, Shigella boydii* and *Shigella dysenteriae* (9). Serological testing further classifies *Shigella* species into more than 55 serotypes through the agglutination reaction of antisera to *Shigella* serotype specific O-antigens (10, 11). Up to 89.6% *Shigella* infections were caused by *S. flexneri* (65.9%) and *S. sonnei* (23.7%) globally (12, 13). The predominant serotype reported in *Shigella* infections has been *S. flexneri* serotype 2a while *S. dysenteriae* serotype 1 has caused the most severe disease (11, 14). Note that for brevity, in all references to *Shigella* serotypes below, *S. sonnei, S. flexneri, S. boydii* and *S. dysenteriae* are abbreviated as SS, SF, SB and SD respectively and a serotype is designated with abbreviated “species” name plus the serotype number e.g. *S. dysenteriae* serotype 1 is abbreviated as SD1.

Enteroinvasive *Escherichia coli* (EIEC) is a pathovar of *E. coli* that causes diarrhoea with less severe symptoms to *Shigella* infections in humans worldwide, particularly in developing countries (8, 13, 15–18). EIEC infections in developed countries are mainly imported (19). EIEC has more than 18 specific E. coli O-serotypes (19, 20). Although the incidence of EIEC is low (17), EIEC serotypes have been associated with outbreaks and sporadic cases of infections (20–22). In contrast to *Shigella*, EIEC infections are not notifiable in many countries (23, 24).

*Shigella* and EIEC have always been considered very closely related and share several characteristics (25–28). *Shigella* and EIEC are both non-motile and lack the ability of ferment lactose (24). Some of EIEC O antigens are identical or similar to *Shigella* O antigens (O112ac, O124, O136, O143, O152 and O164) (26, 29–31). Furthermore, *Shigella* and EIEC both carry the virulence plasmid pINV, which encodes virulence genes required for invasion (32, 33) and contain *ipaH* (invasion plasmid antigen H) genes with the exception of some SB13 isolates (10, 23, 24, 34, 35). *Shigella* and EIEC have arisen from *E. coli* in multiple independent events and should be regarded as a single pathovar of E. coli (25, 26, 28, 36–38). Previous phylogenetic studies suggested that *Shigella* isolates were divided into 3 clusters (C1, C2 and C3) with 5 outliers (SS, SB13, SD1, SD8 and SD10) (25, 38) whereas EIEC isolates were grouped into four clusters (C4, C5, C6 and C7) (26). The seven *Shigella*/EIEC clusters and 5 outliers of *Shigella* are within the broader non-enteroinvasive *E. coli* species except for SB13 which is closer to *Escherichia albertii* (39, 40). WGS-based phylogenomic studies have also defined multiple alternative clusters of *Shigella* and EIEC (23, 28, 41).

The traditional biochemical test for motility and lysine decarboxylase (LDC) activity (42) and molecular test for the presence of *ipaH* gene have been used to differentiate *Shigella* and EIEC from non-enteroinvasive E. coli (24, 43–45). Agglutination with *Shigella*/EIEC associated antiserum further classify *Shigella* or EIEC to serotype level. However, cross-reactivity, strains not producing O antigens, and newly emerged *Shigella* serotypes may all prevent accurate serotyping (10, 46). Serotyping by antigenic agglutination is being replaced by molecular serotyping (47, 48), which can be achieved through examination of the sequences of O antigen biosynthesis and modification genes (8, 24, 49–52).

Recently, PCR-based molecular detection methods targeting the gene *lacY* were developed to distinguish *Shigella* from EIEC (53, 54). However, the ability of the primers described in these methods to accurately differentiate between *Shigella* and EIEC was later questioned (23, 28). With the uptake of whole-genome sequencing technology, several studies have identified phylogenetic clade specific markers, species specific markers and EIEC lineage-specific genes for discrimination between *Shigella* and EIEC and between *Shigella* species (23, 27, 28, 41, 55, 56). More recently, genetic markers *lacY, cadA, Ss_methylase* were used for identification of *Shigella* and EIEC (10). However, these markers failed to discriminate between *Shigella* and EIEC when a larger genetic diversity is considered (23, 28, 55). A Kmer-based approach can identify *Shigella* isolates to the species level but misidentification was also observed (56).

In this study, we aimed to i), identify phylogenetic clusters of *Shigella* and EIEC through large scale examination of publicly available genomes; ii), identify cluster-specific gene markers using comparative genomic analysis of *Shigella* and EIEC accessory genomes for differentiation of *Shigella* and EIEC; iii), develop a pipeline for *Shigella* and EIEC *in silico* serotyping based on the cluster-specific gene markers combined with *Shigella* and EIEC serotype-specific O antigen and H antigen genes. We demonstrate that these cluster-specific gene markers enhance *in silico* serotyping using genomic data. We also developed an automated pipeline for cluster typing and serotyping of *Shigella*/EIEC from WGS data.

## Materials and Methods

### Identification of *Shigella*/EIEC isolates from NCBI database

*E. coli/Shigella* isolates from the NCBI SRA (National Center for Biotechnology Information Sequence Read Archive) as May of 2019 were queried. Raw reads were retrieved from ENA (European Nucleotide Archive). The *ipaH* gene (GenBank accession number M32063.1) was used to screen E. coli/*Shigella* reads using Salmon v0.13.0 (57). Taxonomic classification for *E. coli/Shigella* was confirmed by Kraken v1.1.1 (58). Molecular serotype prediction of *ipaH* negative *Shigella* isolates was performed by ShigaTyper v1.0.6 (10). Isolates that were *ipaH* positive and isolates with designation of SB13 by ShigaTyper were selected as *Shigella*/EIEC database.

The sequence types (STs) and ribosomal STs (rSTs) of *ipaH* gene negative E. coli (non-enteroinvasive E. coli) isolates were examined. STs and rSTs for these isolates were obtained from the E. coli/*Shigella* database in the Enterobase (59) as of May 2019. For STs and rSTs with only one isolate, the isolates were selected. For STs and rSTs with more than one isolates, one representative isolate for each ST and rST were randomly selected. In total, 12,743 ipaH negative E. coli isolates representing 3,800 STs and 11,463 rSTs were selected as non-enteroinvasive E. coli control database.

### Genome sequencing

Whole-genome sequencing (WGS) of 31 EIEC strains used in a previous study (26) was performed by Illumina NextSeq (Illumina, Scoresby, VIC, Australia). DNA libraries were constructed using Nextera XT Sample preparation kit (Illumina Inc., San Diego, CA, USA) and sequenced using the NextSeq sequencer (Illumina Inc.). FASTQ sequences of the strains sequenced in this study were deposited in the NCBI under the BioProject (PRJNA692536).

### Genome assembly and data processing

Raw reads were *de novo* assembled using SPADES v3.14.0 assembler with default settings [http://bioinf.spbau.ru/spades] (60). The metrics of assembled genomes were obtained with QUAST v5.0.0 (61). Three standard deviations (SD) from the mean for contig number, largest contig, total length, GC, N50 and genes were used as quality filter for assembled genomes.

The STs for isolates in *Shigella*/EIEC database was checked by using mlst (https://github.com/tseemann/mlst) with the *E. coli* scheme from PubMLST (62). rSTs were extracted from the E. coli/*Shigella* rMLST database in Enterobase (59) as of May 2019. Serotype prediction for isolates in *Shigella*/EIEC was performed by ShigaTyper v1.0.6 (10). Serotyping of E. coli O and H antigens were predicted by using SerotypeFinder v2.0.1 (63).

### Selection of isolates for *Shigella*/EIEC identification dataset

The selection of isolates for the identification dataset was based on the representative isolates for each ST, rST and serotype of *Shigella* and EIEC in the *Shigella*/EIEC database. For STs and rSTs with only one isolate, the isolate was selected. For STs and rSTs with more than one isolates, one representative isolate for each ST, rST was randomly selected. A representative experimentally confirmed isolate of each serotype of *Shigella* and EIEC was also randomly selected. 72 ECOR strains downloaded from Enterobase (59) and 18 *E. albertii* strains were used as controls for the identification dataset. The details of the identification dataset are listed in Table S1. The remaining isolates in *Shigella*/EIEC database were referred as validation dataset (Table S2).

The identification dataset was used to characterise the phylogenetic relationships of *Shigella* and EIEC. The identification dataset was also used to identify cluster-specific genes. The validation dataset was used to evaluate the performance of cluster-specific gene markers using the *in-silico* serotyping pipeline.

### Phylogeny of *Shigella* and EIEC based on WGS

Three phylogenetic trees including identification tree, confirmation tree and validation tree were constructed by Quicktree v1.3 (64) with default parameters to identify and confirm the phylogenetic clustering of *Shigella* and EIEC isolates. The phylogenetic trees were visualised by Grapetree and ITOL v5 (65, 66).

The identification phylogenetic tree was generated based on isolates in the identification dataset for the characterisation of clusters of *Shigella* and EIEC isolates (Fig. 1). A subset of 485 isolates known to represent each identified cluster from the identification dataset were then selected. The subset of 485 isolates from the identification dataset and 1,872 non-enteroinvasive E. coli isolates from non-enteroinvasive E. coli control dataset (2,357 isolates total) were used to construct a confirmation tree. This tree was used for confirmation of the phylogenetic relationships between identified *Shigella*/EIEC clusters in the identification dataset and non-enteroinvasive E. coli isolates. The validation tree was generated based on 1,159 representative isolates from the validation dataset that were selected in the same way as the identification dataset and a subset of 485 isolates from the identification dataset to assign validation dataset isolates to clusters.

**Figure 1:**
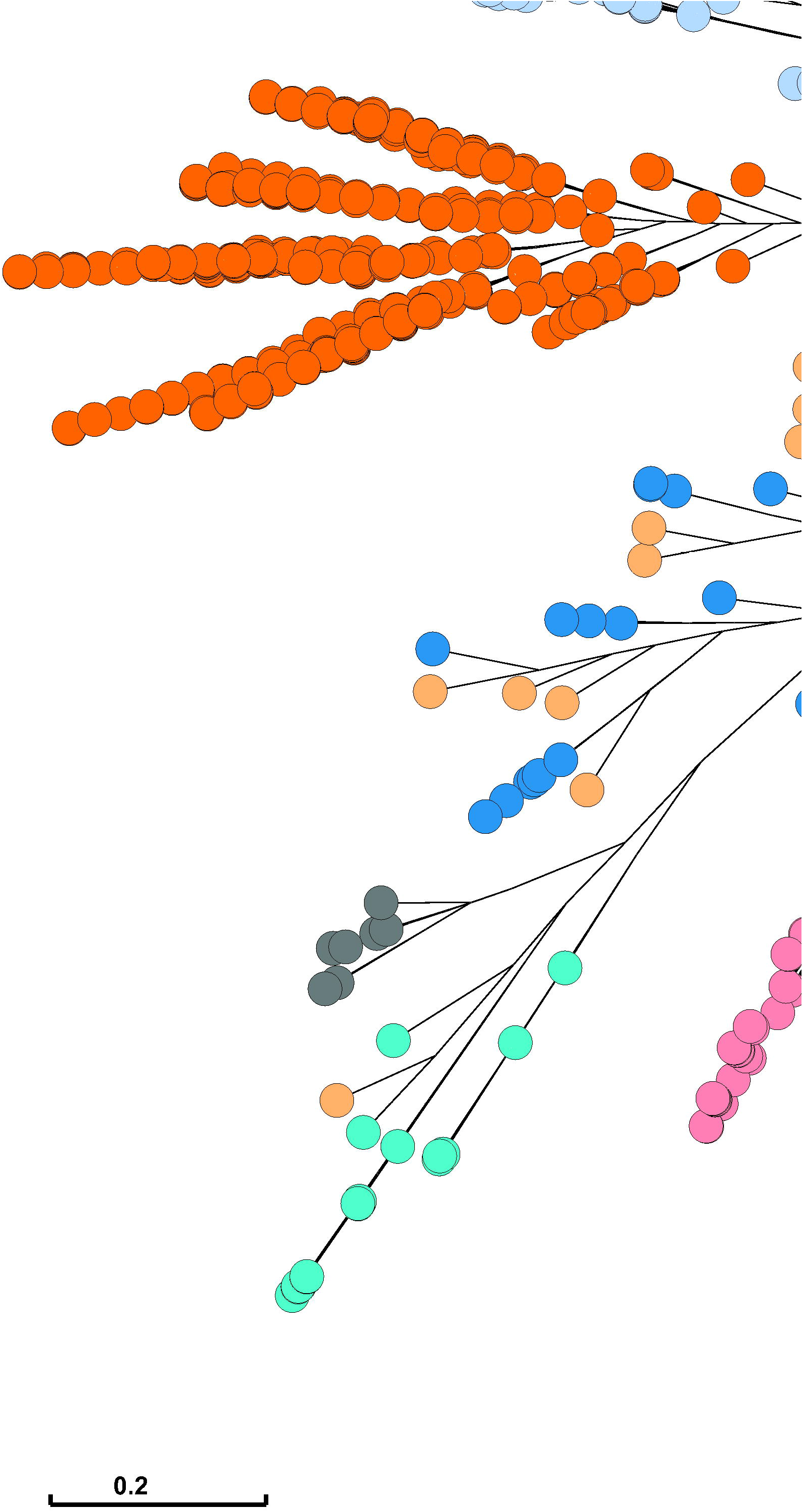
*Shigella*/EIEC cluster Identification phylogenetic tree. Representative isolates from the identification dataset were used to construct the phylogenetic tree by Quicktree v1.3 (64) to identify *Shigella* and EIEC clusters and visualised by Grapetree’s interactive mode. The dendrogram tree shows the phylogenetic relationships of 1879 *Shigella* and EIEC isolates represented in the identification dataset. Branch lengths are log scale for clarity. The tree scales indicated the 0.2 substitutions per locus. *Shigella* and EIEC clusters are coloured. Numbers in square brackets indicate the number of isolates of each identified cluster. CSP is sporadic EIEC lineages.

### Investigation of *Shigella* virulence plasmid pINV

The presence of *Shigella* virulence plasmid pINV in isolates were investigated by using BWA-MEM v0.7.17 (Burrows-Wheeler Aligner) (67) to align isolate raw reads onto the reference sequence of pINV (68) (NC_024996.1). Mapped reads were sorted and indexed using Samtools v1.9 (69). The individual gene coverage from mapping was obtained using Bedtools coverage v2.27.1 (70).

### Identification of the cluster-specific gene markers

Cluster-specific gene markers were identified from *Shigella*/EIEC accessory genomes. The genomes from the identification dataset were annotated using PROKKA v1.13.3 (71). Pan- and core-genomes were analysed by roary v3.12.0 (72) using an 80% sequence identity threshold. The genes specific to each cluster were identified from the accessory genes with an in-house python script. In this study, the number of genomes from a given cluster containing all specific genes for that cluster was termed true positives (TP), the number of genomes from the same cluster lacking any of those same genes was termed false negatives (FN). The number of genomes from other clusters containing all of those same genes was termed false positives (FP).

The sensitivity (True positive rate, TPR) of each cluster-specific gene marker was defined as TP/(TP+FN). The specificity (True negative rate, TNR) was defined as TN/(TN+FP).

### Validation of the cluster-specific gene markers

The ability of cluster-specific gene markers to assign *Shigella*/EIEC isolates was examined by using BLASTN to search against the validation dataset (Table S2) and non-enteroinvasive E. coli control database for the presence of any of the cluster-specific gene marker or a set of cluster-specific gene markers. The BLASTN thresholds were defined as 80% sequence identity and 50% gene length coverage.

### Development of ShigEiFinder, an automated pipeline for molecular serotyping of *Shigella*/EIEC

ShigEiFinder was developed using paired end illumina genome sequencing reads or assembled genomes identify cluster-specific gene markers combined with *Shigella*/EIEC serotype specific O antigen genes (wzx and wzy) and modification genes (Fig. 2, Data S1). We used the same signature O and H sequences from ShigaTyper and SerotypeFinder (Data S2) (10, 63). These include *Shigella* serotype-specifc wzx/wzy genes and modification genes from ShigaTyper and E. coli O antigen and *fliC* (H antigen) genes from SerotypeFinder. *ipaH* gene and 38 virulence genes used in analysis of virulence of 59 sporadic EIEC isolates were also included in the typing reference sequences database. Seven House Keeping (HK) genes *-recA, purA, mdh, icd, gyrB*, *fumC* and *adk* downloaded from NCBI were used for contamination checking.

**Figure 2:**
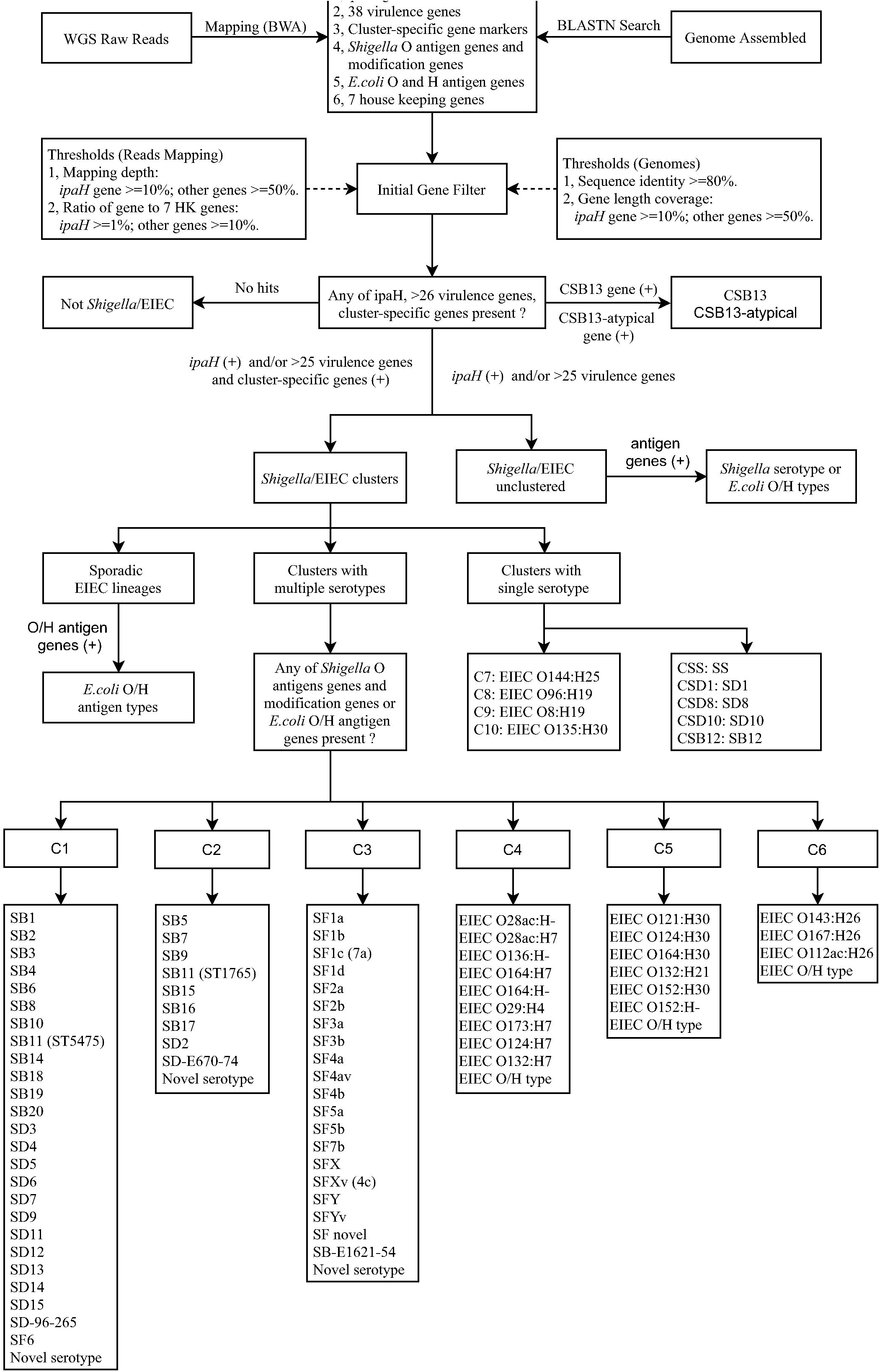
*in silico* serotyping pipeline workflow. Schematic of *in silico* serotyping *Shigella* and EIEC by cluster-specific genes combined with the *ipaH* gene and O antigen and modification genes and H antigen genes, implemented in ShigEiFinder. Both assembled genomes and raw reads are accepted as data input.

Raw reads were aligned to the typing reference sequences by using BWA-MEM v0.7.17 (67). The mapping length percentage and the mean mapping depth for all genes were calculated using Samtools coverage v1.10 (69). To determine whether the genes were present or absent, 50% of mapping length for all cluster-specific genes, virulence genes and O antigen genes and 10% for *ipaH* gene were used as cutoff value. The ratio of mean mapping depth to the mean mapping depth of the 7 HK genes was used to determine a contamination threshold with ratios less than 1% for *ipaH* gene and less than 10% for other genes assigned as contamination. Reads coverage mapped to particular regions of genes were checked by using samtools mpileup v1.10.

Assembled genomes were searched against the typing reference sequences using BLASTN v2.9.0 (73) with 80% sequence identity and 50% gene length coverage for all genes with exception of *ipaH* gene which was defined as 10% gene length coverage.

ShigEiFinder was tested with the identification dataset and validated with the *Shigella*/EIEC validation dataset and non-enteroinvasive E. coli control database. The specificity defined as (1 - the number of non-enteroinvasive E. coli isolates being detected / the total number of non-enteroinvasive E. coli isolates) * 100.

## Results

### Screening sequenced genomes for *Shigella*/EIEC isolates

We first screened available *E. coli* and *Shigella* genomes based on the presence of *ipaH* gene. We examined 122,361 isolates with the species annotation of E. coli (104,256) or *Shigella* (18,105) with paired end illumina sequencing reads available in NCBI SRA database. Of 122,361 isolates, 17,989 isolates were positive to the *ipaH* gene including 455 out of 104,256 E. coli isolates and 17,434 out of 18,105 *Shigella* isolates. The 17,989 *ipaH* positive *E. coli* and *Shigella* genomes and 571 *ipaH* negative “*Shigella*” genomes were checked for taxonomic classification and genome assembly quality. 17,320 *ipaH* positive *E. coli* and *Shigella* genomes and 246 *ipaH* negative “*Shigella”* genomes passed quality filters. Among 246 *ipaH* negative “*Shigella*” genomes, 11 isolates belonged to SB13 by using ShigaTyper (10) while the remaining 235 isolates were classified with taxonomic identifier of E. coli by Kraken v1.1.1 (58) and were removed from analysis. A total of 17,331 genomes including 17,320 *ipaH* positives and 11 SB13 genomes were selected to form the *Shigella*/EIEC database, which contained 429 genomes with species identifier of *E. coli* and 16,902 genomes with species identifier of *Shigella*.

Isolates in *Shigella*/EIEC database were typed using MLST, ShigaTyper and SerotypeFinder. MLST and rMLST divided the 17,331 *Shigella*/EIEC isolates into 252 STs (73 isolates untypeable by MLST) and 1,128 rSTs (3,513 isolates untypeable by rMLST). Of 16,902 genomes with species identifier of *Shigella*, 8,313 isolates and 8,189 isolates were typed as *Shigella* and EIEC respectively by ShigaTyper while 400 isolates were untypeable. ShigaTyper typed the majority of the 8,313 isolates as SF (66.82%) including 25.43% SF2a isolates, followed by SS (19.69%), SB (7.22%) and SD (6.27%).

SerotypeFinder typed 293 of the 429 *E. coli* genomes into 71 *E. coli* O/H antigen types. Among these 293 isolates with typable O/H antigen types, 190 isolates belonged to 22 known EIEC serotypes (O28ac:H-, O28ac:H7, O29:H4, O112ac:H26, O121:H30, O124:H30, O124:H24, O124:H7, O132:H7, O132:H21, O135:H30, O136:H7, O143:H26, O144:H25, O152:H-, O152:H30, O164:H-, O164:H30, O167:H26, O173:H7 and 2 newly emerged EIEC serotypes O96:H19 and O8:H19) (20–22). The remaining 136 of 429 genomes were O antigen untypable and typed to 15 H antigen types only by SerotypeFinder, of which H16 was the predominant type.

### Identification of *Shigella* and EIEC clusters

*Shigella* and EIEC are known to have been derived from *E. coli* independently. To identify previously defined clusters (25, 26) and any new clusters from the 17,331 *Shigella*/EIEC genomes, we selected representative genomes to perform phylogenetic analysis as it was impractical to construct a tree with all genomes. The selection was based on ST, rST and serotype of the 17,331 *Shigella*/EIEC genomes. One isolate was selected to represent each ST, rST and serotype for a total of 1,830 isolates. The selection included 252 STs, 1,128 rSTs, 59 *Shigella* serotypes (21 SB serotypes, 20 SF serotypes, 17 SD serotypes and SS), 22 EIEC known serotypes and 31 other or partial antigen types. A further 31 in-house sequenced EIEC isolates, 18 EIEC isolates used in a previous typing study (41), 72 ECOR strains and 18 *E. albertii* strains were also included to form the identification dataset of 1,969 isolates. Details are listed in Table S1. A phylogenetic tree was constructed based on the identification dataset to identify the clusters (Fig. 1).

All known clusters were identified (Fig. 1) including 3 *Shigella* clusters (C1, C2, C3) and 5 outliers (SD1, SD8, SD10, SB13 and SS) as defined by Pupo et al (25) and 4 EIEC clusters (C4, C5, C6 and C7) defined by Lan et al. (26). Each of these clusters was supported by a bootstrap value of 80% or greater (Fig. S1). 1,789 isolates of the 1,879 *Shigella*/EIEC isolates (1,830 isolates from the *Shigella*/EIEC database, 31 in-house sequenced EIEC isolates and 18 EIEC isolates from Hazen *et al.*) fell within these clusters.

Of the remaining 90 *Shigella*/EIEC unclustered isolates, 31 belonged to 5 *Shigella*/EIEC serotypes including 5 SB13 isolates, 8 SB12 isolates, 2 EIEC O135:H30 isolates, 12 EIEC serotype O96:H19 isolates and 4 EIEC O8:H19 isolates, while 59 isolates were sporadic EIEC isolates which are described in detail in the separate section below. The 5 SB13 isolates were grouped into one lineage within *E. coli* and close to known *Shigella*/EIEC clusters rather than the established SB13 cluster outside *E. coli* which was within the *E. albertii* lineage. The former was previously named as atypical SB13 while the latter was previously named as typical SB13 (39). The 8 SB12 isolates formed one single cluster close to SD1 and atypical SB13 clusters. Two EIEC O135:H30 isolates were grouped as a separate cluster close to C5. Twelve isolates belonging to EIEC serotype O96:H19 and 4 isolates typed as O8:H19 were clustered into two separate clusters, both of which were more closely related to SD8 than other *Shigella*/EIEC clusters. Therefore, atypical SB13 and SB12 were defined as new clusters of *Shigella* while EIEC O96:H19, EIEC O8:H19 and EIEC O135:H30 were defined as C8, C9 and C10 respectively. In total there were 10 *Shigella* clusters and 7 EIEC clusters (Table 1).

**Table 1:** The summary of identified *Shigella*/EIEC clusters and outliers in identification dataset

| Clusters (no of serotypes)<sup>#</sup> | No of isolates | No. STs | No. rSTs | Serotypes |
| --- | --- | --- | --- | --- |
| C1 (25) | 288 | 36 | 166 | SB1-4, SB6, SB8, SB10, SB14, SB18, SB11<sup>b</sup>, SB19-20<sup>b</sup>; SD3-7, SD9, SD11-13, SD14-15<sup>*</sup>, SD-96-265<sup>*</sup>; SF6 |
| C2 (9) | 101 | 19 | 56 | SB5, SB7, SB9, SB11, SB15, SB16, SB17; SD2, SD-E670-74<sup>b</sup>; SD2 |
| C3 (20) | 744 | 81 | 437 | SF1a, SF1b, SF1c (7a), SF2a, SF2b, SF3a, SF3b, SF4a, SF4av, SF4b, SF4bv, SF5a, SF5b, SF7b, SFX, SFXv (4c), SFY, SFYv, SF novel serotype; SB-E1621-54<sup>*</sup> |
| C4 (9) | 51 | 6 | 21 | O28ac:H-, O28ac:H7, O136:H7, O164:H-, O164:H7, O29:H4, O173:H7, O124:H7, O132:H7<sup>*</sup> |
| C5 (6) | 62 | 4 | 15 | O121:H30, O124:H30, O164:H30, O132:H21, O152:H30, O152:H- |
| C6 (3) | 20 | 2 | 6 | O143:H26, O167:H26, O112ac:H26<sup>b</sup> |
| C7 | 10 | 1 | 3 | O144:H25 |
| C8<sup>a</sup> | 12 | 2 | 1 | O96:H19 |
| C9<sup>a</sup> | 4 | 1 | 2 | O8:H19 |
| C10 <sup>#</sup> | 2 | 1 | 1 | O135:H30 |
| CSS | 427 | 39 | 294 |  |
| CSD1 | 70 | 8 | 56 | SD1 |
| CSD8 | 7 | 3 | 3 | SD8 |
| CSD10 | 2 | 2 | 1 | SD10 |
| CSB12<sup>a</sup> | 8 | 2 | 6 | SB12 |
| CSB13 | 7 | 3 | 3 | SB13 |
| CSB13-atypical<sup>a</sup> | 5 | 3 | 3 | SB13 |
| Sporadic EIEC lineages<sup>a</sup> (53) | 59 | 49 | 53 | 53 antigen types |

<sup>#</sup>Numbers in parentheses are the number of serotypes within that cluster.
<sup>a</sup>: Clusters identified as new clusters in this study.
<sup>b</sup>: Serotypes were inconsistent with previous analyses.

### Analysis of the 59 sporadic EIEC isolates

To determine the phylogenetic relationships of the above defined clusters and the remaining 59 sporadic EIEC isolates within the larger non-enteroinvasive *E. coli* population a confirmation tree was generated using 485 isolates representing the known clusters and 1,872 representative non-*Shigella*/EIEC isolates (Fig. S2). The 59 sporadic EIEC isolates including 2 EIEC isolates M2330 (O152:H51) and M2339 (O124:H7) sequenced in this study and 57 isolates were interspersed among non-*Shigella*/EIEC isolates and did not form large clusters. Groups of these isolates that were not previously identified were named as sporadic EIEC lineage followed by their serotype. For example, M2339 (O124:H7) grouped together with one other EIEC isolate with the same O and H antigens O124:H7 and were named ‘sporadic EIEC lineage O124:H7’. There were 53 sporadic EIEC lineages including 5 lineages with 2 or more isolates and 48 lineages with only one isolate. The STs, rSTs and antigen types of these 59 isolates were listed in the Table S1.

Some of the sporadic EIEC isolates fell into STs containing *ipaH* negative isolates. We therefore examined the presence of the pINV virulence plasmid in the sporadic EIEC isolates. We selected 38 genes that are essential for virulence including 35 genes (12 *mxi* genes, 9 *spa* genes, 5 *ipaA-J* genes, 6 *ipgA-F* genes as well as *acp, virB, icsB*) in the conserved entry region encoding the Mxi-Spa-Ipa type III secretion system and its effectors and 3 regulator genes (*virF, virA* and *icsA/virG*) (24, 33, 68) and determined the presence of pINV in the 59 sporadic EIEC isolates by mapping the sequence reads onto a pINV reference sequence (68). Reads from 18 *non-Shigella/EIEC* isolates that shared the same ST as one of 58 sporadic isolates were positive for these genes.

The number of essential virulence genes with mapped reads in the 59 sporadic EIEC isolates were analysed (Fig. S3). Those isolates containing more than 25 of the 38 essential virulence genes were defined as virulence plasmid positive. While isolates containing between 13 and 25 were defined as intermediate and less than 13 were defined as virulence plasmid negative.

The 2 newly sequenced sporadic EIEC isolates (M2330 and M2339) were positive for the virulence plasmid and of the other 57 sporadic EIEC isolates, 39 isolates were positive, 9 isolates were negative and 9 isolates were intermediate (Table S1). The results were compared with those *non-Shigella/EIEC* isolates belonging to the same ST. The virulence plasmid was absent in all *non-Shigella/EIEC* isolates while all sporadic EIEC isolates in these STs were either positive or intermediate. Therefore, this analysis confirmed the sporadic isolates belonged to EIEC and the STs contained both EIEC and non-EIEC isolates.

### Identification of cluster-specific gene markers

In this study, cluster-specific gene markers were either a single gene present in all isolates of a cluster and absent in all other isolates or a set of genes (two or more) that as a combination were only found in one cluster. For the marker sets, a subset of cluster-specific gene markers for a given cluster could be found in other clusters but the entire set was only found in the target cluster.

Comparative genomic analysis on 1,969 accessory genomes from the identification dataset was used to identify cluster-specific gene markers or marker sets. Multiple candidate cluster-specific gene markers or marker sets of markers for each of 17 *Shigella*/EIEC clusters and 53 sporadic EIEC lineages were identified through screening the accessory genes from 1,969 genomes. These gene markers or marker sets were 100% sensitive to clusters but with varying specificity. The cluster-specific gene markers or marker sets with the lowest FP rates were then selected from candidate cluster-specific gene markers by BLASTN searches against genomes in the identification dataset using 80% sequence identity and 50% gene length threshold.

Five single cluster-specific gene markers (C7, C10, SB12, SB13 and atypical SB13) and 12 sets of cluster-specific gene markers (C1, C2, C3, C4, C5, C6, C8, C9, SS, SD1, SD8 and SD10) were selected for *Shigella*/EIEC cluster typing. The sensitivity and specificity for each cluster-specific gene marker or a set of cluster-specific gene markers for the identification dataset were listed in Table 2. The cluster-specific gene markers or marker sets were all 100% sensitive and 100% specific with the exception of C1 (99.94% specificity), C3 (99.91% specificity) and SS (99.8% specificity). A single specific gene for each of 53 sporadic EIEC lineages were also selected with the exception of one lineage which has a set of 2 genes. These genes were all 100% sensitive and specific for a given sporadic EIEC lineage.

**Table 2:** The sensitivity and specificity of cluster-specific genes

| Clusters | Cluster-specific<br>genes (Single/sets)<sup>b</sup> | Identification dataset (1969 isolates) |  |  |
| --- | --- | --- | --- | --- |
|  |  | No of isolates | Sensitivity | Specificity |
| C1 | Set of 4 genes | 288 | 100 | 99.94<sup>a</sup> |
| C2 | Set of 3 genes | 101 | 100 | 100 |
| C3 | Set of 3 genes | 744 | 100 | 99.59<sup>a</sup> |
| C4 | Set of 2 genes | 51 | 100 | 100 |
| C5 | Set of 3 genes | 62 | 100 | 100 |
| C6 | Set of 2 genes | 20 | 100 | 100 |
| C7 | Single gene | 10 | 100 | 100 |
| C8 | Set of 2 genes | 12 | 100 | 100 |
| C9 | Set of 2 genes | 4 | 100 | 100 |
| C10 | Single gene | 2 | 100 | 100 |
| CSS | Set of 5 genes | 427 | 100 | 99.87<sup>a</sup> |
| CSD1 | Set of 2 genes | 70 | 100 | 100 |
| CSD8 | Single gene | 7 | 100 | 100 |
| CSD10 | Single gene | 2 | 100 | 100 |
| CSB12 | Single gene | 8 | 100 | 100 |
| CSB13 | Single gene | 7 | 100 | 100 |
| CSB13-atypical | Single gene | 5 | 100 | 100 |
| 53 Sporadic EIEC lineages | Single gene / lineage | 59 | 100 | 100 |

<sup>a</sup>:The specificity of cluster-specific gene set less than 100% was due to at least one FP found in that set.
<sup>b</sup>: The sequences of these genes were listed in Data S1.

All cluster-specific gene markers, 37 in total (5 single, 32 genes in 12 sets) and 54 sporadic EIEC lineages specific gene markers were located on chromosome but one of C4 gene markers and 5 sporadic EIEC lineages specific genes were located on plasmid by NCBI BLAST searching. None of the cluster-specific gene markers were contiguous in the genomes. The location of these cluster-specific gene markers was determined by BLASTN against representative complete genomes of *Shigella*/EIEC containing gene features downloaded from NCBI GenBank. In those cluster or sporadic lineages with no representative complete genome specific gene markers were named using their cluster or sporadic EIEC lineage followed by the cluster or lineage number. For example, C7 specific gene marker was named “C7 specific gene”.

The functional characterization of these specific gene markers were identified from RAST annotation (74). For 37 cluster-specific gene markers, 22 had known functions and 15 encoded hypothetical proteins with unknown functions, while 11 sporadic EIEC lineages specific gene markers were identified with known functions and 43 were hypothetical proteins with unknown functions. The location and functions of specific gene markers are listed in Table S3.

### Validation of cluster-specific gene markers

The ability of cluster-specific gene markers to correctly assign *Shigella*/EIEC isolates was evaluate with 15,501 *Shigella*/EIEC isolates in the validation dataset, 12,743 isolates from non-enteroinvasive *E. coli* control database.

Using cluster-specific gene markers, 15,443 of the 15,501 (99.63%) *Shigella*/EIEC isolates were assigned to clusters which included 15,337 *Shigella* isolates, 102 EIEC isolates, 4 sporadic EIEC isolates, and 38 (0.24%) isolates with more than one clusters. Twenty of the 15,501 (0.13%) *Shigella*/EIEC isolates were not assigned to any of identified clusters.

To confirm the assignment of cluster-specific gene markers, we constructed a “validation” phylogenetic tree (Fig. S4) using 1,159 representative isolates from the validation dataset and a subset of 485 isolates from each cluster from the identification dataset. Isolates that grouped with known cluster isolates (from identification dataset) with strong bootstrap support were assigned to that cluster. All 1,159 isolates were grouped into known clusters on the validation phylogenetic tree. The cluster-specific gene markers assignments were entirely consistent with cluster assignments by phylogenetic tree.

We tested cluster-specific gene markers with the 12,743 non-enteroinvasive *E. coli* isolates. The *Shigella*/EIEC cluster-specific gene markers were highly specific with specificity varying from 98.8% to 100% for cluster-specific genes and 97.02% to 100% for sporadic EIEC specific genes. Details are listed in Table S4.

### Development of an automated pipeline for molecular serotyping of *Shigella*/EIEC

Above results showed that cluster-specific gene markers were sensitive and specific and can distinguish *Shigella* and EIEC isolates. We therefore used these genes combined with established *Shigella*/EIEC serotype specific O antigen and H antigen genes to develop an automated pipeline for *in silico* serotyping of *Shigella*/EIEC (Fig. 2).

The pipeline is named *Shigella* EIEC Cluster Enhanced Serotype Finder (ShigEiFinder). ShigEiFinder can process either paired end Illumina sequencing reads or assembled genomes (installable package: https://github.com/LanLab/ShigEiFinder, online too: https://mgtdb.unsw.edu.au/ShigEiFinder/). ShigEiFinder classifies isolates into Non-*Shigella*/EIEC*, Shigella* or EIEC clusters based on the presence of *ipaH* gene, number of virulence genes, cluster specific genes. The “Not *Shigella*/EIEC” assignment was determined by the absence of the *ipaH* gene, virulence genes (<26) and absence of cluster-specific gene markers. The “*Shigella* or EIEC clusters” assignments were made based on the presence of *ipaH* gene, and/or more than 25 virulence genes together with the presence of any of cluster-specific gene markers or marker set, whereas the presence of *ipaH* gene and/or more than 25 virulence genes with absence of any of cluster-specific gene markers were assigned as “*Shigella*/EIEC unclustered”.

*Shigella* and EIEC isolates were differentiated and serotypes were assigned after cluster assignment. ShigEiFinder predicts a serotype through examining the presence of any of established *Shigella* serotype specific O antigen and modification genes and E. coli O and H antigen genes that differentiate the serotypes as ShigaTyper and SerotypeFinder (10, 63). A “novel serotype” is assigned if no match to known serotypes.

Two pairs of *Shigella* serotypes, SB1/SB20 and SB6/SB10, are known to be difficult to differentiate as they share identical O antigen genes (10, 46, 75). ShigaTyper used a heparinase gene for the differentiation of SB20 from SB1 and *wbaM* gene for the separation of SB6 from SB10. We found that fragments of the heparinase and *wbaM* genes may be present in other serotypes and cannot accurately differentiate SB1/SB20 and SB6/SB10. We found a SB20 specific gene which encoded hypothetical proteins with unknown functions and located on a plasmid by comparative genomic analysis of all isolates in C1 accessory genome. The SB20 specific gene can reliably differentiate SB20 from SB1and also one SNP each in *wzx* and *wzy* genes that can differentiate SB6 from SB10. We used these differences (Data S1) in ShigEiFinder for the prediction of these serotypes.

### The accuracy and specificity of ShigEiFinder in cluster typing

The accuracy of ShigEiFinder was tested with 1,969 isolates (1,969 assembled genomes and 1,951 Illumina reads [note no reads available for 18 EIEC isolates from NCBI) from the identification dataset and 15,501 isolates from the validation dataset. The results are listed in Table 3.

**Table 3:**
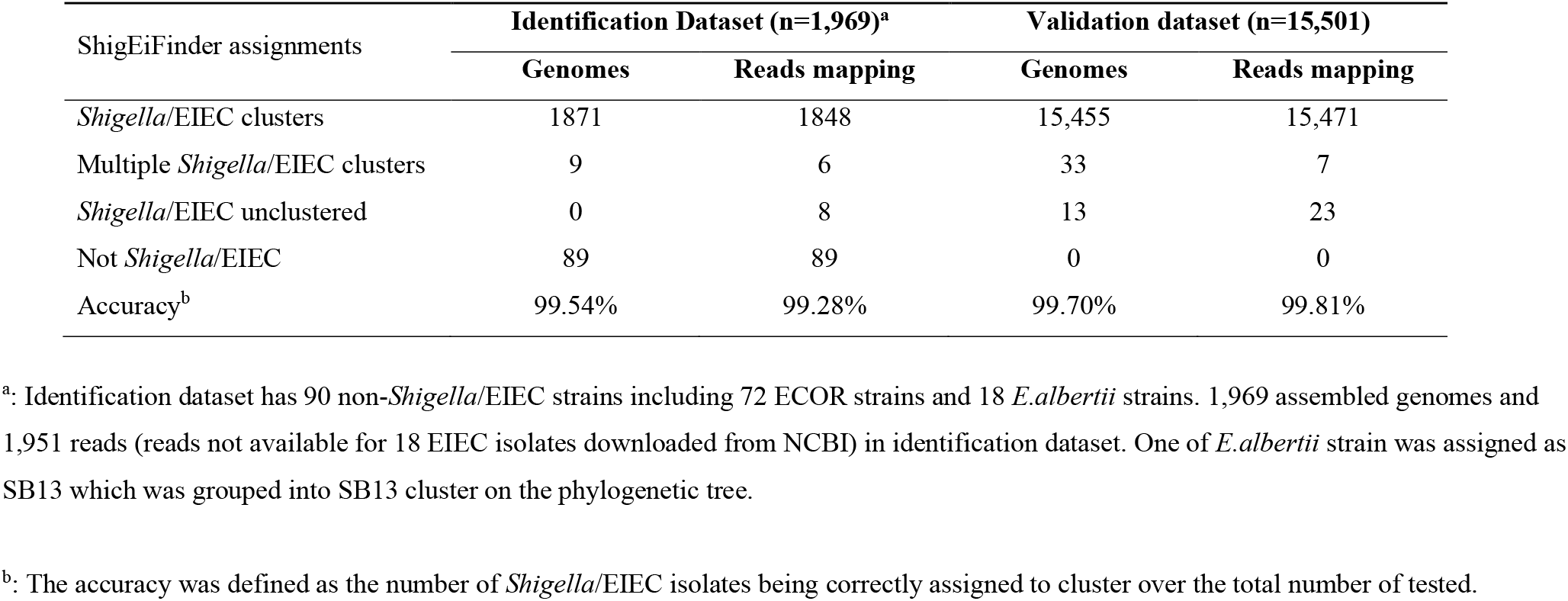
The accuracy of ShigEiFinder with identification dataset and validation dataset

ShigEiFinder was able to assign 99.54% and 99.28% of the isolates in the identification dataset to clusters for assembled genomes and read mapping respectively. The accuracy was 99.70% and 99.81% for assembled genomes and read mapping respectively when applied to the validation dataset. Discrepancies were observed between assembled genomes and read mapping (Table 3). There were more isolates assigned to *“Shigella/EIEC* unclustered” in read mapping, in contrast there were more isolates assigned to multiple clusters in genome assemblies. The specificity of ShigEiFinder was 99.40% for assembled genomes and 99.38% for read mapping when evaluated with 12,743 non-*Shigella*/EIEC *E. coli* isolates. An additional 2 isolates were detected as sporadic EIEC lineages by read mapping.

### Comparison of ShigEiFinder and ShigaTyper

To demonstrate ShigEiFinder for differentiation of *Shigella* from EIEC and enhancement of cluster based serotyping, the comparison of read mapping results between ShigEiFinder and the existing *in silico Shigella* identification pipeline ShigaTyper (10) was performed with 488 isolates used in ShigaTyper and 15,501 isolates from *Shigella*/EIEC validation dataset used in the present study.

The 488 isolates used in ShigaTyper consisted of 23 other species, 45 *E. coli* isolates and 420 *Shigella* isolates. ShigEiFinder identified 23 other species isolates and 453 out of 465 *E. coli* and *Shigella* isolates, in agreement with ShigaTyper assignment. ShigEiFinder also assigned the remaining 3 EIEC isolates and 9 (either multiple *wzx* or no *wzx* genes found) isolates untypeable by ShigaTyper to *Shigella*/EIEC clusters.

ShigEiFinder assigned 15,471 of 15,501 *Shigella*/EIEC isolates to *Shigella* or EIEC clusters and then to a serotype. The accuracy of ShigEiFinder to correctly assign isolates to *Shigella* or EIEC clusters was 99.81% (15,471/15,501). By contrast, ShigaTyper assigned 7,277 isolates (46.95%) to *Shigella*, 7.976 isolates (51.45%) to EIEC, 177 (1.14%) isolates to multiple *wzx* genes and failed to type 71 (0.46%) isolates.

The predicted serotype of 7,277 (46.96%) *Shigella* isolates by ShigaTyper agreed with the results of ShigEiFinder. For 8,224 isolates typed as EIEC or untypable by ShigaTyper, 99.73% (8,202/8,224) of the isolates were assigned to *Shigella* or EIEC clusters by ShigEiFinder (Table 4). Of these isolates, the majority belonged to SS, SD1 and SF which were erroneously predicted as EIEC by ShigaTyper.

**Table 4:**
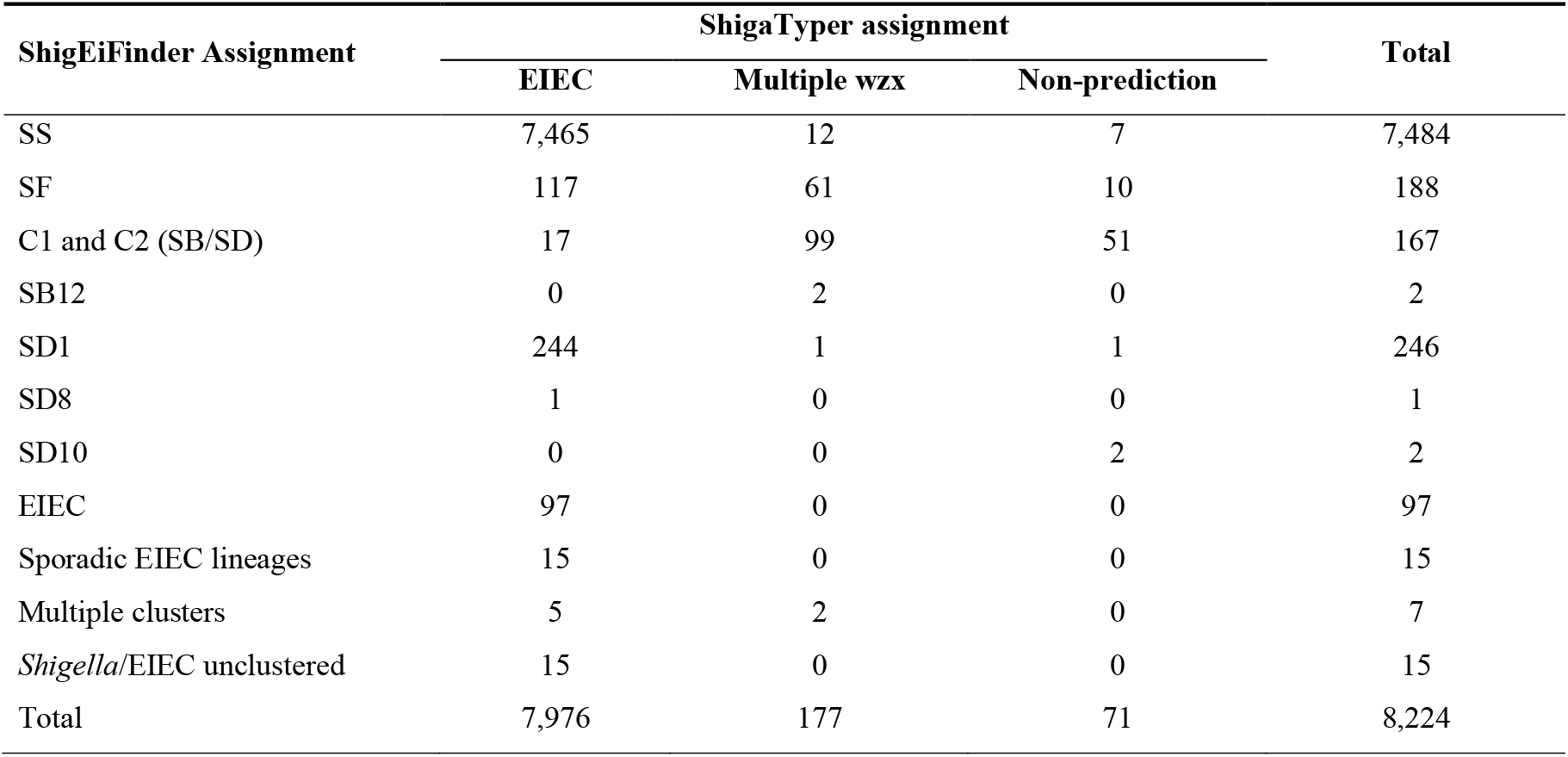
Discrepant assignment of 8,224 isolates by ShigEiFinder and Shigatyper

## Discussion

*Shigella* and EIEC cause human bacillary dysentery with similar invasion mechanisms, however the pathogenicity of these 2 groups varies (8, 43). The prevalence of each of the four *Shigella* “species” also varies (11–13). Differentiation of *Shigella* and EIEC from each other is important for epidemiologic and diagnostic investigations. However, their similar physiological, biochemical and genetic characteristics make this differentiation difficult.

### Determining phylogenetic clusters for better separation of *Shigella* isolates from EIEC

From a phylogenetic perspective, *Shigella* and EIEC strains consisted of multiple phylogenetic lineages derived from commensal *E. coli*, which do not reflect the nomenclature of *Shigella* and EIEC (23, 25, 26, 28, 38, 41). In the present study, we identified all phylogenetic clusters of *Shigella* and EIEC through large scale examination of publicly available genomes. Phylogenetic results demonstrated that *Shigella* isolates had at least 10 clusters while EIEC isolates had at least 7 clusters. The 10 *Shigella* clusters included the 7 previously defined lineages including 3 major clusters (C1, C2 and C3) and 5 outliers (SD1, SD8, SD10, SB13 and SS) (25) and 2 newly identified clusters (SB12 and SB13-atypical). The 7 EIEC clusters consisted of 4 previously defined EIEC clusters (C4, C5, C6 and C7) (26) and 3 newly identified EIEC clusters (C8 EIEC O96:H19, C9 EIEC O8:H19 and C10 EIEC O135:H30).

Our WGS-based phylogeny provided high resolution for assigning *Shigella* and EIEC isolates to clusters. Several serotypes that are currently increasing in frequency (SB19, SB20, SD14, SD15, SD provisional serotype 96-626) (76–79) were assigned to clusters and five new clusters/outliers were identified. SB13 isolates in this study formed two known lineages. One lineage was located outside of *Shigella*/EIEC clusters and represented the outlier SB13 which is in fact belonging to the newly defined species *E. albertii* (25, 26, 38, 39). The second lineage was with *E. coli*, and was defined as atypical SB13 previously (39). The newly identified *Shigella* outlier SB12 was previously grouped into C3 based on housekeeping gene trees (25, 38) but was seen as outliers in two other studies (28, 56).

Newly identified clusters C8 (EIEC O96:H19) and C9 (EIEC O8:H19) represented the emergence of novel EIEC serotypes. A recent study revealed that EIEC serotype O96:H19 (C8) could be the result of a recent acquisition of the invasion plasmid by commensal E. coli (80). The EIEC serotype O8:H19 (C9) had not been reported previously.

Apart from the 17 major and outlier clusters of *Shigella* and EIEC, the presence of 53 sporadic EIEC lineages indicated greater genetic diversity than has been observed previously. Isolates belonging to these sporadic EIEC groups were more closely related to non-enteroinvasive E. coli isolates than to major *Shigella*/EIEC lineages. However, 41 of these isolates, representing 38 sporadic EIEC lineages, carried pINV. *Shigella* and EIEC both carry the *Shigella* virulence plasmid pINV which is vital for virulence and distinguishes *Shigella*/EIEC from other *E. coli* (24, 33, 68). Therefore, these isolates may represent recently formed EIEC lineages through acquisition of the pINV. The remaining 18 isolates contained the *ipaH* gene but may or may not carry pINV. It is possible that these strains carried very low copy number of pINV or the pINV plasmid was lost during culture.

### Highly sensitive and specific cluster-specific gene markers for differentiation of *Shigella* and EIEC isolates

Several studies have identified phylogenetic related genomic markers for discrimination of *Shigella* and EIEC and between *Shigella* species (23, 27, 28, 41, 55, 56). However, these phylogenetic analyses were performed only with a small number of genomes (23, 28, 55). In addition, non-invasive E. coli isolates were included in some of the phylogenetic clusters identified (28) which led to non-invasive E. coli isolates being identified by the markers.

We identified cluster-specific gene markers for each respective cluster which were only composed of *Shigella* or EIEC isolates. Sets of cluster-specific gene markers were identified for those clusters where no single suitable marker is present. The combination of genes enhances the specificity of cluster-specific gene markers as demonstrated by the 100% sensitivity and very high specificity in this analysis (Table 2). Genes specific to each of the 53 sporadic EIEC lineages were also identified and they were sensitive and specific, although it should be noted that these values are based on very small sample sizes.

The cluster-specific gene markers or marker sets can be used to differentiate *Shigella*/EIEC from non-enteroinvasive *E. coli* independent of *ipaH* gene. The *ipaH* gene as a molecular target has been used to differentiate *Shigella* and EIEC from non-enteroinvasive E. coli (24, 43–45). In our study, the cluster-specific gene markers were specific to *Shigella*/EIEC with 98.8% to 100% specificity when evaluated on non-enteroinvasive *E. coli* control database, providing confidence that the cluster-specific genes or sets are robust markers for the identification of *Shigella*/EIEC. 53 sporadic EIEC lineage specific gene markers also have very high specificity (97.02% to 100%) against non-enteroinvasive *E. coli* control database.

The cluster-specific gene markers or marker sets are able to assign *Shigella*/EIEC isolates to correct clusters in 99.63% of cases and can clearly distinguish *Shigella* isolates from EIEC when applied to the validation dataset. While ShigaTyper assigned 46.95% isolates to *Shigella* and 51.45% isolates to EIEC in the same dataset we tested, leading to a large proportion of isolates incorrectly assigned. The majority of the isolates predicted as EIEC by ShigaTyper were SS or SD1 as they belonged to SS and SD1 specific STs and were positive to a set of SS or SD1 specific gene markers and grouped into SS or SD1 cluster on our phylogenetic tree. The genes used in ShigaTyper were SS specific marker Ss_methylase gene (81, 82) together with SS O antigen wzx gene. However, SS specific marker Ss_methylase gene was found in other *Shigella* serotypes and EIEC (10) and SS O antigen wzx gene were located on a plasmid which is frequently lost (83). Similarly, the SD1 O antigen genes used in ShigaTyper were plasmid-borne which may also lead to inconsistent detection (84, 85). A previous study identified 6 loci to distinguish EIEC from *Shigella* (23). We searched the 6 loci against our *Shigella*/EIEC database and found that some *Shigella* isolates were misidentified as EIEC isolates, such as SD8 isolates incorrectly identified as EIEC subtype 13. Our cluster-specific genes can differentiate SD8 isolates from EIEC with 100% accuracy. Therefore, the cluster-specific gene markers marker sets described here provided nearly perfect differentiation of *Shigella* from EIEC.

The cluster-specific gene markers or marker sets are able to differentiate SS and SF (with exception of SF6) from SB and SD. SF and SS are the major cause of *Shigella* infections, accounting for up to 89.6% annual cases (11–13). Differentiation of SS and SF isolates from SB and SD is also beneficial for diagnosis and surveillance. A recent study identified “species” specific markers for the detection of each of the four *Shigella* “species” and validated with only one isolate per species (55). A molecular algorithm based on *Shigella* O antigen genes can detect 85% of SF isolates (52). In contrast, a set of SF specific genes in our study can correctly identify SF isolates with 99.62% accuracy.

The cluster-specific gene markers or marker sets can also assign *Shigella*/EIEC isolates to serotype level if the cluster has single serotype such as SD1, SD8, SD10, SB13, SB12, EIEC O144:H25 (C7), EIEC O96:H19 (C8), EIEC O8:H19 (C9) and EIEC O135:H30 (C10). The remaining EIEC, SF, SB and SD serotypes were distributed over the major clusters C4-6, C3, C1 and C2 respectively. Cluster-specific gene markers or marker sets combined with serotype associated O antigen and modification genes can further identify these isolates to serotype level if the cluster has multiple serotypes. Once an isolate is assigned to a cluster, only serotype associated O antigen and modification genes found in that cluster need be examined. This allows the elimination of ambiguous or incorrect serotype assignments that may otherwise occur, increasing the overall accuracy of the method.

### Cluster-specific gene marker based ShigEiFinder can accurately type *Shigella* and EIEC

To facilitate the use of cluster-specific gene markers or marker sets for typing, we developed an automated pipeline, ShigEiFinder, for *in silico* molecular serotyping of *Shigella/EIEC.* ShigEiFinder provided *Shigella/* EIEC differentiation as well as serotype prediction by yielding “presence or absence” of cluster-specific gene markers or marker sets combined with *Shigella*/EIEC O antigen genes and modification genes in a query isolate (either reads or assembled genomes). We showed 99.70% and 99.81% accuracy to assign isolates to the correct clusters from 15,501 *Shigella*/EIEC isolates in validation dataset for the assembled genomes and reads mapping respectively. In contrast, the existing *in silico Shigella* serotyping pipeline ShigaTyper had 46.95% accuracy for reads mapping when tested with the same validation dataset, with 51.45% of isolates in validation dataset being predicted as EIEC by ShigaTyper.

The genetic determinants used in ShigaTyper for differentiation of *Shigella* from EIEC and identification of SS were *lacY, cadA, Ss_methylase*, SS and SD1 O antigen *wzx* genes (10). As discussed above some of these genes were found to be non-specific in this study. Compared with ShigaTyper, the cluster-specific gene markers used in ShigEiFinder for identification of *Shigella* and EIEC provided higher discriminatory power than ShigaTyper. ShigEiFinder also provided a high specificity with 99.40% for assembled genomes and 99.38% for reads mapping.

ShigEiFinder can differentiate *Shigella* isolates from EIEC and distinguish SS and SF (with exception of SF6) isolates from SB and SD accurately. It also can identify SD1 isolates directly. ShigEiFinder was able to serotype over 59 *Shigella* serotypes and 22 EIEC serotypes. Therefore, ShigEiFinder will be useful for clinical, epidemiological and diagnostic investigations and the cluster-specific gene markers identified could be adapted for metagenomics or culture independent typing.

## Conclusion

This study analysed over 17,000 publicly available *Shigella*/EIEC genomes and identified 10 clusters of *Shigella*, 7 clusters of EIEC and 53 sporadic types of EIEC. Cluster-specific gene markers or marker sets for the 17 major clusters and 53 sporadic types were identified and found to be valuable for *in silico* typing. We additionally developed ShigEiFinder, a freely available *in silico* serotyping pipeline incorporating the cluster-specific gene markers to facilitate serotyping of *Shigella*/EIEC isolates using genome sequences with very high specificity and sensitivity.

## Authors and contributors

Conceptualization: R.L, M.P.; Investigation: X.Z., M.P., T.N., S.K.; Methodology: M.P., R.L. Writing – original draft: X.Z.; Writing – review and editing: M.P., R.L.

## Acknowledgements

The authors thank Duncan Smith and Robin Heron from UNSW Research Technology Services for computing assistance.

## Data bibliography

Zhang X, Payne M, Nguyen T, Kaur S, Lan R. All the sequencing data generated within this study, NCBI BioProject number (PRJNA692536).

## Abbreviations

SS: *Shigella sonnei*
SF: *Shigella flexneri*
SB: *Shigella boydii*
SD: *Shigella dysenteriae*
EIEC: Enteroinvasive *Escherichia coli*
NCBI SRA: National Center for Biotechnology Information Sequence Read Archive
ST: sequence type
rST: ribosomal ST
MLST: Multilocus sequence typing
rMLST: Ribosomal MLST
ECOR: *Escherichia coli* reference collection
WGS: wholegenome sequencing
TP: true positive
FN: false negative
FP: false positive
HK: House Keeping

## Supplementary Material

**Figure S1: Identification phylogenetic tree**

An identification phylogenetic tree constructed by Quicktree v1.3 (64) and visualised by ITOL v5 shows the phylogenetic relationships of 1879 *Shigella* and EIEC isolates in identification dataset. The tree scales indicated the 0.01 substitutions per locus. *Shigella* and EIEC clusters are colored. The internal branches are colored to represent the bootstrap values. Green color indicates the maximum bootstrap value (1). The red color shows the minimum bootstrap value (0). Each of cluster is well supported by bootstrap value. CSP is sporadic lineages.

**Figure S2-A: Confirmation phylogenetic tree**

A confirmation phylogenetic tree was constructed by Quicktree v1.3 (64) based on 2375 isolates and visualised by Grapetree’s interactive mode. The tree shows the phylogenetic relationships between identified *Shigella*/EIEC clusters in identification dataset and non-enteroinvasive *E.coli* isolates. Branch lengths are log scale for clarity. The tree scales indicated the 0.1 substitutions per locus. Known *Shigella* and EIEC clusters from identification dataset are colored. Numbers in square brackets indicate the number of isolates of each identified cluster. CSP is sporadic lineages.

**Figure S2-B: Confirmation phylogenetic tree**

A confirmation phylogenetic tree constructed by Quicktree v1.3 (64) and visualised by ITOL v5 shows the phylogenetic relationships between identified *Shigella*/EIEC clusters in identification dataset and non-enteroinvasive *E.coli* isolates. The tree scales indicated the 0.01 substitutions per locus. *Shigella* and EIEC clusters are colored. The internal branches are colored to represent the bootstrap values. Green color indicates the maximum bootstrap value (1). The red color shows the minimum bootstrap value (0). Each of cluster is well supported by bootstrap value. CSP is sporadic lineages.

**Figure S3: Distribution of mapped 38 virulence genes in 59 sporadic isolates**

The presence of *Shigella* virulence plasmid pINV in 59 sporadic isolates in identification dataset was determined by the mapped 38 virulence genes. Detailed genes were described in Results “**Investigation of *Shigella* virulence plasmid pINV in 59 sporadic isolates**”. Three categories were defined based on the number of virulence genes mapped to isolate. Virulence plasmid positive: > 25 genes mapped to isolate; Intermediate: 13 to 25 genes mapped to isolate; Virulence plasmid negative: less than 13 genes mapped to isolate.

**Figure S4 (A): Validation phylogenetic tree**

A validation tree was generated by Quicktree v1.3 (64) and visualised by Grapetree’s interactive mode to assign representative isolates in validation dataset to clusters. Branch lengths are log scale for clarity. The tree scales indicated the 0.2 substitutions per locus. Known *Shigella* and EIEC clusters from identification dataset are colored. Numbers in square brackets indicate the number of isolates of each identified cluster. Isolates in validation dataset are colored white. The isolates are assigned to clusters if they grouped into known cluster isolates. CSP is sporadic lineages.

**Figure S4 (B): Validation phylogenetic tree**

A validation phylogenetic tree was constructed by Quicktree v1.3 (64) and visualised by ITOL v5 to assign representative isolates in validation dataset to clusters. The tree scales indicated the 0.01 substitutions per locus. *Shigella* and EIEC clusters are colored. The internal branches are colored to represent the bootstrap values. Green color indicates the maximum bootstrap value (1). The red color shows the minimum bootstrap value (0). Each of cluster is well supported by bootstrap value. Isolates that grouped with known cluster isolates (from identification dataset) with strong bootstrap support are categorised into that cluster. CSP is sporadic lineages.

**Table S1**: 1,969 isolates used in identification dataset

**Table S2**: 15,501 isolates used in validation dataset

**Table S3**: The location and function of cluster-specific genes

**Table S4**: The results of cluster-specific gene markers tested with 12,743 non-enteroinvasive *E.coli* isolates

**Data S1**: Algorithms incorporated into the ShigEiFinder

**Data S2**: Genetic signature O and H genes from ShigaTyper and SerotypeFinder

## Data Availability Statement

Custom python scripts used in this study are available from the authors on request.

