## Supplemental Data S1 for "Cluster-specific gene markers enhance *Shigella* and Enteroinvasive *Escherichia coli in silico* serotyping"

**Algorithms incorporated into the ShigEiFinder**

ShigEiFinder stands for *Shigella* EIEC Cluster Enhanced Serotype Finder and is a cluster-specific gene marker based *in silico* pipeline developed for differentiation of *Shigella* and enteroinvasive *E. coli* (EIEC) and serotyping of *Shigella* and EIEC. ShigEiFinder is available as a web tool and on github.

Note that for brevity, in all references to *Shigella* serotypes below, *Shigella sonnei*, *Shigella flexneri*, *Shigella boydii* and *Shigella dysenteriae* are abbreviated as SS, SF, SB and SD respectively and a serotype is designated with abbreviated “species” name plus the serotype number e.g. *Shigella dysenteriae* serotype 1 is abbreviated as SD1.

**Typing reference sequences used in ShigEiFinder**

The typing reference sequences consisted of cluster-specific gene markers and sporadic EIEC lineages specific gene markers from this study, *ipaH* gene, 38 virulence genes, *Shigella* serotype specific O antigen genes collected from ShigatTyper (1), *E.coli* O antigen genes and *fliC* genes collected from SerotypeFinder (2) and 7 House Keeping (HK) genes from the MLST (3) scheme.

The cluster-specific gene markers and markers sets and sporadic EIEC lineages specific gene markers are listed as supplementary material with file name in Table S3. The 38 virulence genes are listed in “Analysis of the 59 sporadic EIEC isolates” section in the main text. *Shigella* and *E.coli* O and H antigen genes are listed as supplementary material with file name in Data S2.

All sequences are listed in fasta format available at <https://github.com/LanLab/ShigEiFinder>.

**ShigEiFinder input**

Either paired end Illumina sequencing reads or assembled genomes are acceptable.

**ShigEiFinder output**

ShigEiFinder output included the sample, presence of *ipaH* gene, number of virulence genes, cluster assignment, serotype, *E. coli* O and H antigen present and any further notes for the result in a tabular format.

**Decisions made for genes present or absent**

The presence or absence of genes were determined by the cutoff value of gene length coverage for assembled genomes and the mapping length percentage and the ratio of mean mapping depth to the average mean mapping depth of 7 HK genes (Table 1). For example, the *ipaH* gene was defined as present if mapping length coverage was over 10% together with the ratio of mean depth to the average mean depth of 7 HK genes was over 1% from reads mapping.

**Table 1: Thresholds used for determination of genes present or absent**

| Typing reference genes | Genomes | Reads mapping | |
| --- | --- | --- | --- |
|  | Gene length coverage | Mapping length coverage | Ratio to  7 HK |
| *ipaH* gene | 10% | 10% | 1% |
| Vriulence genes | 50% | 50% | 10% |
| Cluster-specfic gene markers | 50% | 50% | 10% |
| O antigen and H antigen genes | 50% | 50% | 10% |

**Decisions made for cluster assignment and serotyping**

The *Shigella* or EIEC cluster assignment was determined by the presence of cluster-specific gene markers that were only found within a single *Shigella*/EIEC cluster. Where markers set (more than one gene) was used to identify a cluster, all genes must be present for a cluster to be called. ShigEiFinder also used 38 virulence genes from the pINV invasive plasmid to determine whether the plasmid was present in the genome. When more than 25 of these genes were present, the isolate was considered to be pINV positive.

The presence of cluster-specific gene markers or markers sets combined with the presence of *ipaH* gene and/or virulence genes the isolate was assigned to *Shigella* or EIEC cluster (Table 2).

The isolate assigned as *Shigella*/EIEC unclustered could be any new cluster that can not be detected by any of cluster-specific gene markers or markers set. *Shigella*/EIEC unclustered isolate could also be the presence of all genes in markers set except one of the genes from markers set which had mapping ratio between 1% and 10%, therefore making it unclustered (11 isolates of 15,501 isolates in validation dataset).

**Table 2: The cluster-specific gene markers based cluster assignment**

| Cluster assignment | *ipaH* gene | >=26 virulence genes | cluster-specific gene/set |
| --- | --- | --- | --- |
| *Shigella*/EIEC clusters | + | +/- | + |
| *Shigella*/EIEC unclustered | + | +/- | - |
| SB13/SB13-atypical | - | - | + |
| Not *Shigella*/EIEC | - | - | - |

“+”: gene presence; “+/-”: can be present or absent; “-”: gene absence.

The serotype then assigned based on the presence of *Shigella* serotype specific O antigen genes and modification genes or *E. coli* O and H antigen genes. A “novel serotype” is assigned if no match to known serotypes.

**Low level contamination check and note for *Shigella*/EIEC unclustered**

The gene markers with mapping ratio between 1% and 10% demonstrated that the genes in the genomes may not be sequenced very well or a potential contamination. ShigEiFinder noted as “Possible contamination by *Shigella*/EEIC strain or low cluster-specific gene mapping depth to HK genes in cluster X”.

The genes may have mapping ratio between 1% and 10% are listed in Table 3.

**Table 3: Gene markers with mapping ratio between 1% and 10%**

| Gene markers | Number of isolates of 15,501 isolates |
| --- | --- |
| C1_gene_2 | 8 |
| C1_gene_4 | 1 |
| C5_gene_1 | 1 |
| CSS_gene_3 | 1 |

**Additional subsets of gene markers used for *Shigella*/EIEC clusters assignment**

To increase the accuracy of typing, we added additional subsets of genes to eliminate the known false presences for cluster-specific gene markers. For example, the combination of C1 specific markers set and CSB12 specific gene marker can eliminate false presence isolate of CSB12 from C1, if both genes were presence, the isolate could be assigned CSB12 while if CSB12 specific gene was absent, the isolate was assigned as C1. There were 6 subsets of combined genes incorporated into the ShigEiFinder for elimination of false presences (Table 4).

**Table 4: Subsets of combined gene markers for elimination of false cluster assignment**

| Subset 1 | C1 markers set | CSB12 gene | Cluster Assignment |
| --- | --- | --- | --- |
| Isolate | + | + | CSB12 |
| Isolate | + | - | C1 |
| Isolate | - | + | CSB12 |
| Subset 2 | C1 markers set | CSD1 markers set | Cluster Assignment |
| Isolate | + | + | CSD1 |
| Isolate | + | - | C1 |
| Isolate | - | + | CSD1 |
| Subset 3 | C1 markers set | C2 markers set | Cluster Assignment |
| Isolate | + | + | C2 |
| Isolate | + | - | C1 |
| Isolate | - | + | C2 |
| Subset 4 | C3 markers set | C5 markers set | Cluster Assignment |
| Isolate | + | + | C3 |
| Isolate | + | - | C3 |
| Isolate | - | + | C5 |
| Subset 5 | C5 markers set | C8 markers set | Cluster Assignment |
| Isolate | + | + | C8 |
| Isolate | + | - | C5 |
| Isolate | - | + | C8 |
| Subset 6 | C2 markers set | CSS markers set | Cluster Assignment |
| Isolate | + | + | C2 |
| Isolate | + | - | C2 |
| Isolate | - | + | CSS |

“+”: gene presence; “-”: gene absence.

**Serotyping SB1/SB20 within C1**

SB1 and SB20 share identical O antigen genes. For better differentiation of SB1 from SB20, we analysed C1 subbranch on identification tree (Fig.1 in main text). The 21 isolates with presence of SB1 wzx and wzy genes were grouped into one subbranch which consisted of 2 lineages, lineage I and lineage II as Table 5.

**Table 5: The distribution of SB1/SB20 isolates in two lineages**

| Lineages | ShigaTyper assignment | | | | |
| --- | --- | --- | --- | --- | --- |
|  | SB1 | SB20 | EIEC | Untypeable | Total |
| Lineage I | 11 | 0 | 1 | 2^a^ | 14 |
| Lineage II | 4 | 2 | 2 | 1 | 9^b^ |

^a^: One isolate with the presence of heparinase gene which was used in ShigaTyper to separate SB20 form SB1.

^b^: All 9 isolates had heparinase gene either full length or fragments by BLASTN search.

To determine phylogenetic level of the 2 lineages, HierBAPS (4) analysis was performed and the results indicated that 2 lineages were at the different phylogenetic level. Therefore lineage I was defined as potential SB1 lineage and lineage II was defined as SB20 lineage (Figure below). Based on phylogenetic analysis, we identified an SB20 specific gene by comparing 288 accessory genomes in C1 from the identification dataset. The gene was validated with *Shigella*/EIEC validation dataset C1 isolates. The isolate was assigned as SB20 with the presence of SB20 specific gene and SB1 wzx/wzy genes, otherwise the isolate was SB1 with the only SB1 wzx/wzy genes present.

**Figure: Subbranch of C1 on identification tree**

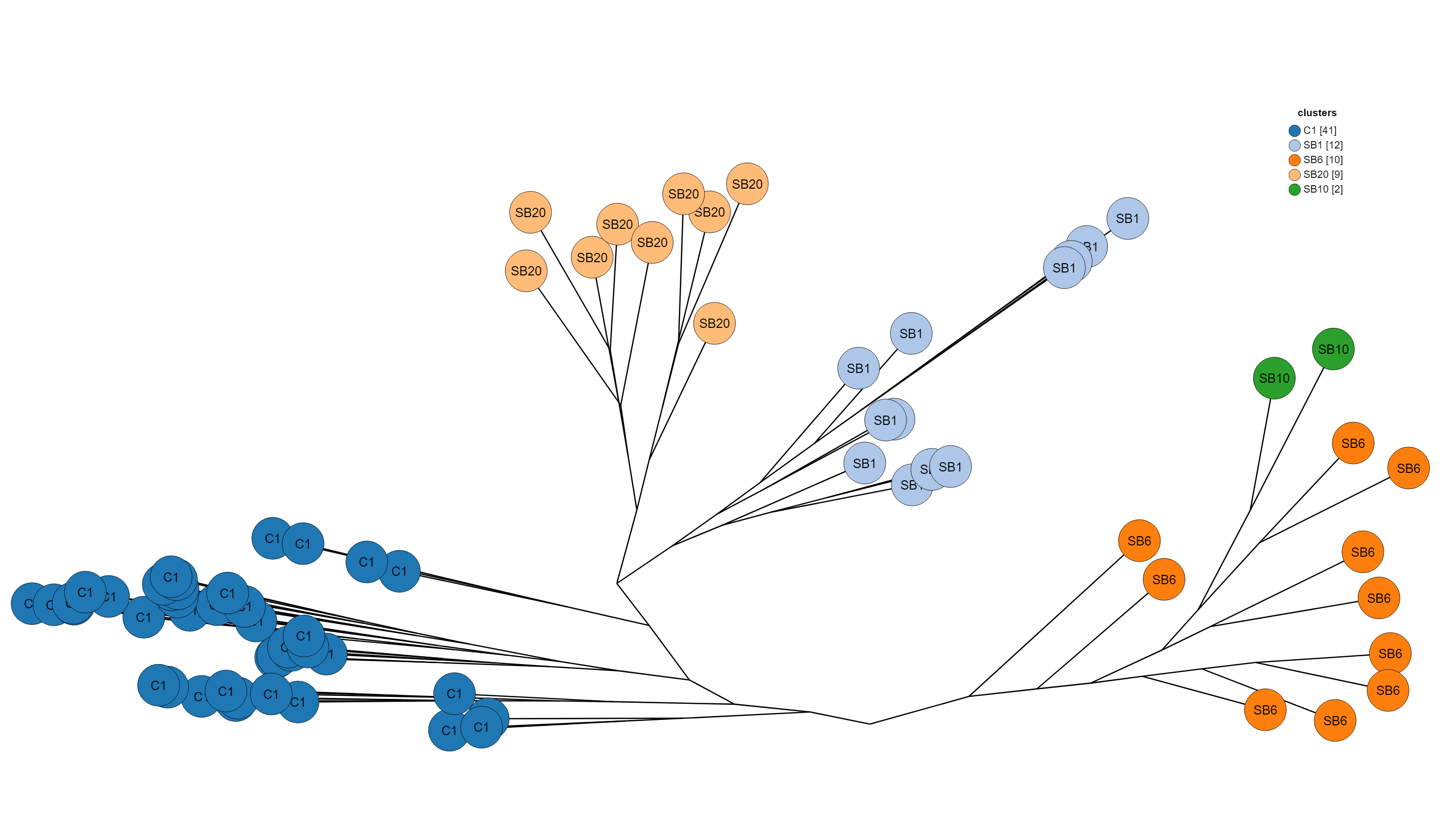

**Serotyping SB6/SB10 within C1**

SB6 and SB10 share identical O antigen genes as well. The SNP in SB10 wzx and SB10 wzy genes at positions 904 and 141 respectively were used to separate SB6 from SB10. For assembled genomes, we first checked the SNP positions that were covered in the blast search with 100% identify for SB10. The isolate was classified as SB10 if the SB10 SNPs were present. Otherwise the isolate was assigned as SB6. Samtools mpileups was used to gather the nucleotide base at the SNP positions for reads mapping. The isolate was SB10 if the SB10-SNPs was found. The absence of SNP was assigned as SB6.

**Serotyping EIEC O164/O124**

The *E.coli* O164 and O124 O antigen genes are near identical with > 99.4% identity (5). There was a 2-base indel at positions 429 and 430 in *wfep* gene for O164. We used this indel to differentiate O164 from O124. The isolate was assigned as O164 if the indel was found.

**Multiple H antigens**

There are multiple variants for one type of H antigen. To solve the multiple H antigen types assigned, the highest match was chosen as the H antigen present.

**SF serotyping within C3**

C3 has all SF serotypes except for SF6 which is grouped into C1. We used the established SF O antigen genes and modification genes including *gtr*, *oac* and *opt* genes to type SF within C3 (Table 6). ShigEiFInder assigned all possibilities when there was a multiple match of combinations of modification genes. The isolate was classified as SFY if there was only backbone O antigen genes present. While the isolate was assigned as SF novel serotype with no genes found and the note was given with the presence or absence of genes.

**Table 6: The combination of O antigen genes and modification genes used for SF serotyping**

|  | *wzx_1-5_* | *wzx_6_* | *gtrI* | *gtrIC* | *gtrII* | *gtrIV* | *gtrV* | *gtrX* | *oac* | *oac1b* | *oacB* | *oacC* | *oacD* | *optII* | *optIII* |
| --- | --- | --- | --- | --- | --- | --- | --- | --- | --- | --- | --- | --- | --- | --- | --- |
| SF1a | + | - | + | - | - | - | - | - | - | - | +/- | - | - | - | - |
| SF1b | + | - | + | - | - | - | - | - | - | + | +/- | - | - | - | - |
| SF1c(7a) | + | - | + | + | - | - | - | - | - | - | - | - | - | - | - |
| SF1d | + | - | + | - | - | - | - | + | - | - | - | - | - | - | - |
| SF2a | + | - | - | - | + | - | - | - | - | - | +/- | - | +/- | - | - |
| SF2b | + | - | - | - | + | - | - | + | - | - | - | - | +/- | - | - |
| SF3a | + | - | - | - | - | - | - | + | +/- | - | - | - | +/- | - | - |
| SF3b | + | - | - | - | - | - | - | - | + | - | - | - | - | - | - |
| SF4a | + | - | - | - | - | + | - | - | - | - | - | - | - | - | - |
| SF4av | + | - | - | - | - | + | - | - | - | - | - | - | - | - | + |
| SF4b | + | - | - | - | - | + | - | - | + | - | - | - | - | - | - |
| SF5a | + | - | - | - | - | - | + | - | - | - | + | - | - | - | - |
| SF5b | + | - | - | - | - | - | + | + | - | - | - | - | - | - | - |
| SF7b | + | - | - | + | - | - | - | - | + | - | - | - | - | - | - |
| SFX | + | - | - | - | - | - | - | + | - | - | - | - | +/- | - | - |
| SFXv(4c) | + | - | - | - | - | - | - | + | - | - | - | - | +/- | + | - |
| SFY | + | - | - | - | - | - | - | - | - | - | +/- | - | +/- | - | - |
| SFYv | + | - | - | - | - | - | - | - | - | - | - | - | + | + | + |
| SF6 | - | + | - | - | - | - | - | - | - | - | - | +/- | - | - | - |

“+”: gene presence and highlighted in pink color.

“+/-”: can be present or absent.

“-”: gene absence.
