## Supplemental Data S2 for "Cluster-specific gene markers enhance *Shigella* and Enteroinvasive *Escherichia coli in silico* serotyping"

Data S2: ***Shigella*/EIEC serotypes specific O and H antigens used in ShigEiFinder**

*Shigella* serotype specific O antigen genes were collected from ShigaTyper (1). *E.coli* O antigen genes and *fliC* genes were collected from SerotypeFinder (2).

| Sequences | Accession number |
| --- | --- |
| SD1_*wzx*, SD1_*wzy* | L07293 |
| SD1-rfp | CP000640 |
| SD2_*wzx*, SD2_*wzy* | EU296404 |
| SD3_*wzx*, SD3_*wzy* | EU296415 |
| SD4_*wzx*, SD4_*wzy* | EU296402 |
| SD5_*wzx*, SD5_*wzy* | EU294174 |
| SD6_*wzx*, SD6_*wzy* | EU296414 |
| SD7_*wzx*, SD7_*wzy* | AY380835 |
| SD8_*wzx*, SD8_*wzy* | EU294166 |
| SD9_*wzx*, SD9_*wzy* | EU296416 |
| SD10_*wzx*, SD10_*wzy* | EU294178 |
| SD11_*wzx*, SD11_*wzy* | EU294172 |
| SD12_*wzx*, SD12_*wzy* | EU294169 |
| SD13_*wzx*, SD13_*wzy* | EU294167 |
| SD14_*wzx*, SD14_*wzy* | CP026832 |
| SD15_*wzx*, SD15_*wzy* | CP026834 |
| SDP 96-265_*wzx*, *wzy* | CP026819 |
| SDP E670-74_*wzx*,*wzy* | CP027027 |
| SB1_*wzx*, SB1_*wzy* | AY630255 |
| SB2_*wzx*, SB2_*wzy* | EU296418 |
| SB3_*wzx*, SB3_*wzy* | EU296407 |
| SB4_*wzx*, SB4_*wzy* | AF402312 |
| SB5_*wzx*, SB5_*wzy* | AF402313 |
| SB6_*wzx*, SB6_*wzy* | AF402314 |
| SB7_*wzx*, SB7_*wzy* | EU296411 |
| SB8_*wzx*, SB8_*wzy* | EU294163 |
| SB9_*wzx*, SB9_*wzy* | AF402315 |
| SB10_*wzx*, SB10_*wzy* | AY693427 |
| WbaM | AY693427 |
| SB11_*wzx*, SB11_*wzy* | AY529126 |
| SB12_*wzx*, SB12_*wzy* | EU296406 |
| SB13_*wzx*, SB13_*wzy* | AY369140 |
| SB14_*wzx*, SB14_*wzy* | EU296409 |
| SB15_*wzx*, SB15_*wzy* | EU296412 |
| SB16_*wzx*, SB16_*wzy* | DQ371800 |
| SB17_*wzx*, SB17_*wzy* | DQ875941 |
| SB18_*wzx*, SB18_*wzy* | AY948196 |
| SB19_*wzx*, SB19_*wzy* | CP026814 |
| Heparinase | CP016036 |
| SBP E1621-54_*wzx*,*wzy* | CP026810 |
| SF *wzx*_1-5_ gene | AE005674 |
| SF6 *wzx* gene | EU294165 |
| SF *gtrI* | AF139596 |
| SF *gtrIC* | FJ905303 |
| SF *gtrII* | AF021347 |
| SF *gtrIV* | AF288197 |
| SF *gtrV* | U82619 |
| SF *gtrX* | L05001 |
| SF *oacA* | AF547987 |
| SF *oac1b* | JN377795 |
| SF *oacB* | NC_004337 (SF0315) |
| SF *oacC* | AKMW01000058 |
| SF *oacD* | NC_004337 (SF0309) |
| SF *optIII* | KC020049 |
| SF Xv *optII* | CP001385 (SFxv_5135) |
| SS_*wzx*, SS_*wzy* | AF285971 |
| O1_*wzx*, O1_*wzy* | GU299791 |
| O2­_*wzx*, O2_*wzy* | EU549863 |
| O4­_*wzx*, O4_*wzy* | AY568960 |
| O6­_*wzx*, O6_*wzy* | AJ426045 |
| O7_*wzx*, O7_*wzy* | AF125322 |
| O8­_*wzx*, O8_*wzy* | AF013583 |
| O8­_*wzm*, O8_*wzt* | AB010150 |
| O12_*wzx*, O12_*wzy* | AB811600 |
| O13_*wzx*, O13_*wzy* | EU296422 |
| O16­_*wzx*, O16_*wzy* | AB811601 |
| O17­_*wzx*, O17_*wzy* | AB812084 |
| O18­ac_*wzx*, O18ac_*wzy* | AB811603 |
| O21­_*wzx*, O21_*wzy* | EU694098 |
| O22­_*wzx*, O22_*wzy* | AB811606 |
| O25­_*wzx*, O25_*wzy* | GU014554 |
| O26­_*wzx*, O26_*wzy* | AF529080 |
| O28ac­_*wzx*, O28ac_*wzy* | DQ462205 |
| O29_*wzx*, O29_*wzy* | EU294173 |
| O32_*wzx*, O32_*wzy* | EU296410 |
| O36_*wzx*, O36_*wzy* | AB811613 |
| O39­_*wzx*, O39_*wzy* | AB811616 |
| O40_*wzx*, O40_*wzy* | EU296417 |
| O50­_*wzx*, O50_*wzy* | AB811624 |
| O53­_*wzx*, O53_*wzy* | EU289392 |
| O71_*wzx*, O71_*wzy* | GU445927 |
| O77_*wzx*, O77_*wzy* | AB972416 |
| O79_*wzx*, O79_*wzy* | EU294162 |
| O86_*wzx*, O86_*wzy* | AY220982 |
| O89_*wzx*, O89_*wzy* | AB812038 |
| O92_*wzx*, O92_*wzy* | AB812040 |
| O93_*wzx*, O93_*wzy* | AB812041 |
| O96_*wzx*, O96_*wzy* | AB812043 |
| O102_*wzx*, O102_*wzy* | JX087966 |
| O105_*wzx*, O105_*wzy* | EU294171 |
| O110_*wzx*, O110_*wzy* | AB812049 |
| O111_*wzx*, O111_*wzy* | JN887675 |
| O112ab_*wzx*, O112ab _*wzy* | EU296413 |
| O112ac_*wzx*, O112ac_*wzy* | EU296405 |
| O117_*wzx*, O117_*wzy* | EU694096 |
| O118_*wzx*, O118_*wzy* | HM204927 |
| O121_*wzx*, O121_*wzy* | JN859209 |
| O124_*wzx*, O124_*wzy* | EU296419 |
| O129_*wzx*, O129_*wzy* | EU296424 |
| O130_*wzx*, O130_*wzy* | EU296421 |
| O132_*wzx*, O132_*wzy* | AB812056 |
| O135_*wzx*, O135_*wzy* | EU296423 |
| O136_*wzx*, O136_*wzy* | AB812059 |
| O143_*wzx*, O143_*wzy* | EU294164 |
| O144_*wzx*, O144_*wzy* | AB812062 |
| O147_*wzx*, O147_*wzy* | DQ868766 |
| O148_*wzx*, O148_*wzy* | DQ167407 |
| O149_*wzx*, O149_*wzy* | DQ868764 |
| O151_*wzx*, O151_*wzy* | HM204926 |
| O152_*wzx*, O152_*wzy* | EU294170 |
| O155_*wzx*, O155_*wzy* | AY657020 |
| O162_*wzx*, O162_*wzy* | AB812067 |
| O164_*wzx*, O164_*wzy* | EU296420 |
| O167_*wzx*, O167_*wzy* | EU296408 |
| O173_*wzx*, O173_*wzy* | GU068046 |
| O180_*wzx*, O180_*wzy* | JQ751058 |
| O183_*wzx*, O183_*wzy* | AB627352 |
| H1_*fliC* | AB028471 |
| H2_*fliC* | AIHA01000023 |
| H4_*fliC* | AJ605764 |
| H4_*fliC* | AJ605765 |
| H4_*fliC* | AJ536600 |
| H5_*fliC* | AY249990 |
| H5_*fliC* | AY337469 |
| H6_*fliC* | AIEY01000041 |
| H7_*fliC* | AY337468 |
| H7_*fliC* | AKML01000326 |
| H7_*fliC* | ANLT01000257 |
| H7_*fliC* | ANLJ01000383 |
| H7_*fliC* | AOES01000098 |
| H7_*fliC* | AMVH01000352 |
| H7_*fliC* | AF228487 |
| H7_*fliC* | AF228496 |
| H7_*fliC* | AF228495 |
| H7_*fliC* | AB334575 |
| H7_*fliC* | AB334574 |
| H7_*fliC* | AF228494 |
| H7_*fliC* | AF228491 |
| H7_*fliC* | AF228492 |
| H7_*fliC* | AB028474 |
| H8_*fliC* | AJ865465 |
| H9_*fliC* | AY249994 |
| H10_*fliC* | AF169320 |
| H11_*fliC* | AY337472 |
| H12_*fliC* | AY337471 |
| H14_*fliC* | AY249998 |
| H16_*fliC* | AB128919 |
| H16_*fliC* | JH954529 |
| H16_*fliC* | JH953794 |
| H16_*fliC* | AY337476 |
| H16_*fliC* | AY337477 |
| H16_*fliC* | AY337475 AY2500001 |
| H16_*fliC* | AY250000 |
| H18_*fliC* | AY250001 |
| H19_*fliC* | AY337479 |
| H19_*fliC* | AY250002 |
| H20_*fliC* | AY250003 |
| H21_*fliC* | AIHL01000060 |
| H24_*fliC* | K72 (H25w) |
| H25_*fliC* | AGSG01000116 |
| H26_*fliC* | AY250008 |
| H26_*fliC* | AY337483 |
| H27_*fliC* | AY250009 |
| H28_*fliC* | AY250010 |
| H30_*fliC* | AY250011 |
| H30_*fliC* | AY337483 |
| H31_*fliC* | AY250013 |
| H33_*fliC* | AY250015 |
| H40_*fliC* | AJ884568 |
| H42_*fliC* | AY250021 |
| H45_*fliC* | AY250023 |
| H48_*fliC* | AY250025 |
| H49_*fliC* | AY250026 |
| H51_*fliC* | AY250027 |
