## Supplementary figures and images for "Cluster-specific gene markers enhance *Shigella* and Enteroinvasive *Escherichia coli in silico* serotyping"

### Figure S1.tiff

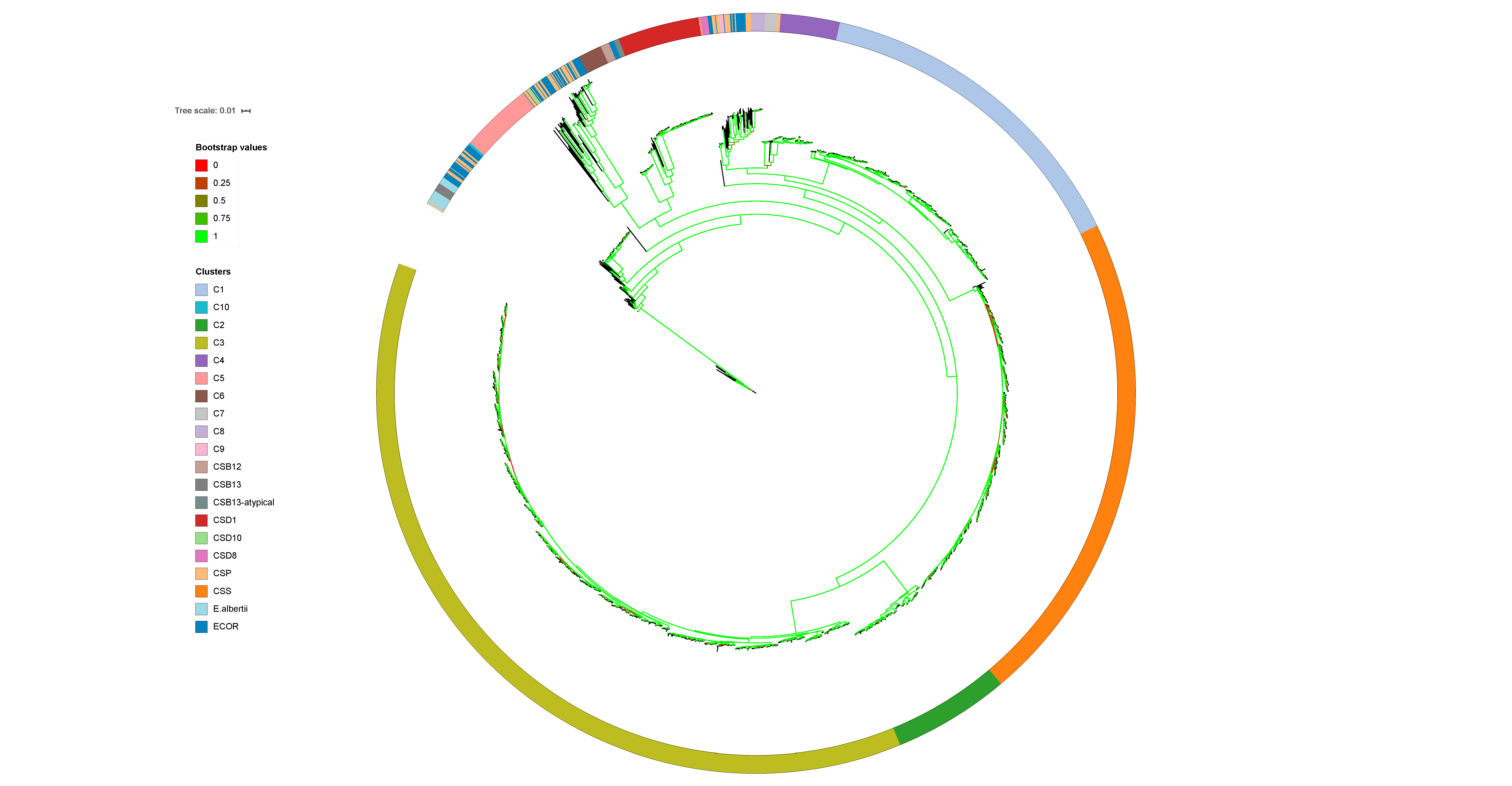

### Figure S2A.tiff

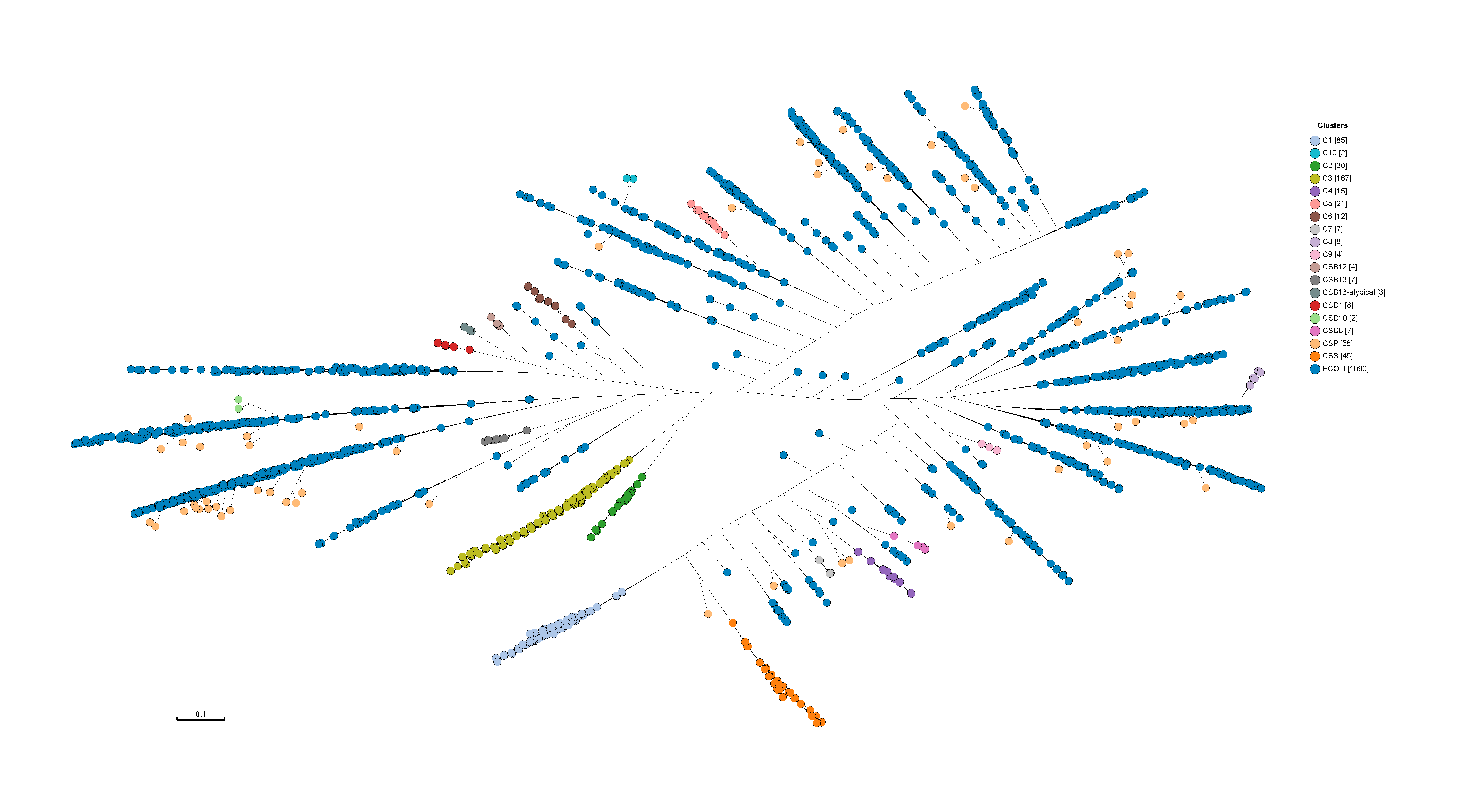

### Figure S2B.tiff

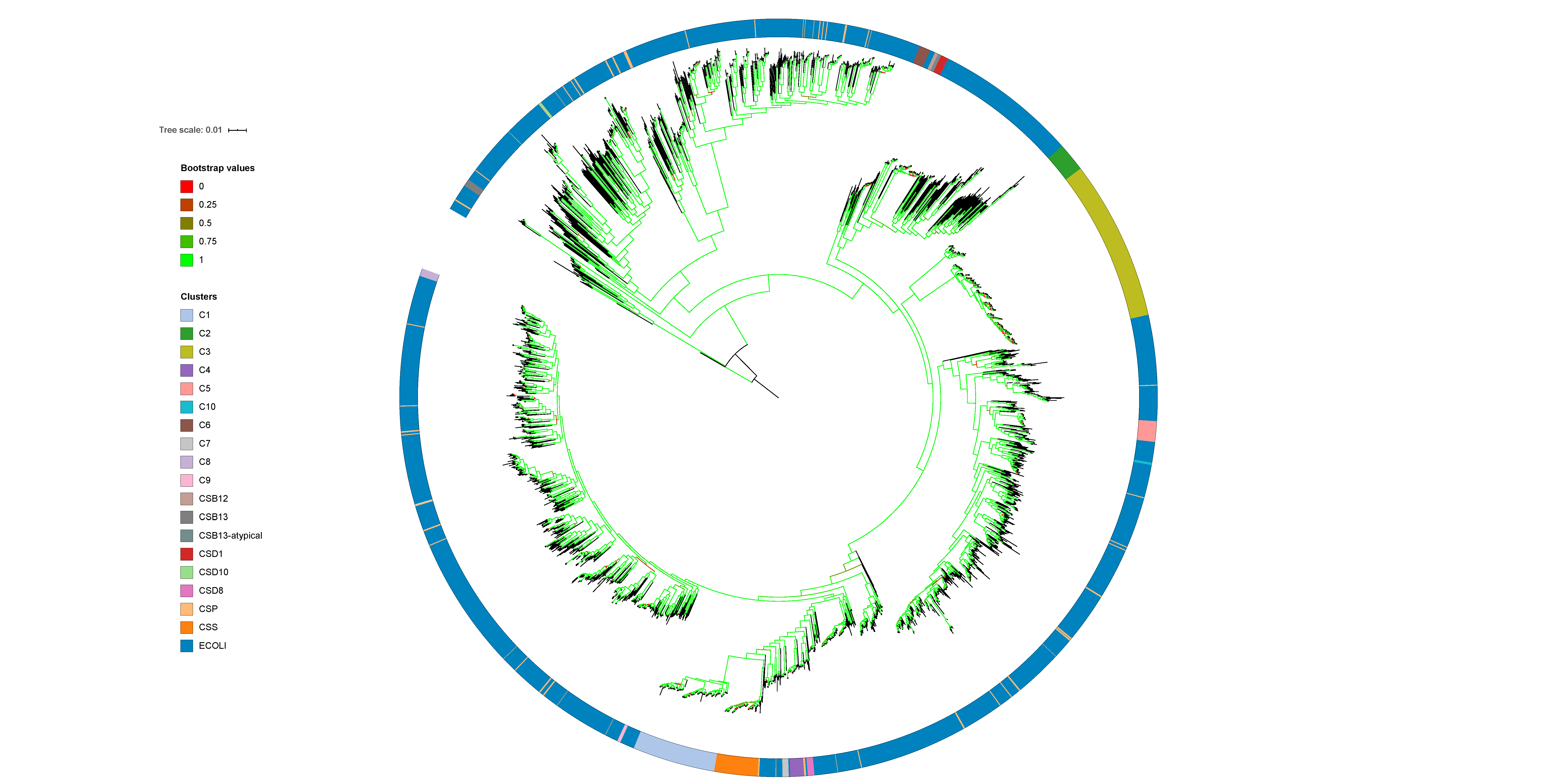

### Figure S3.tiff

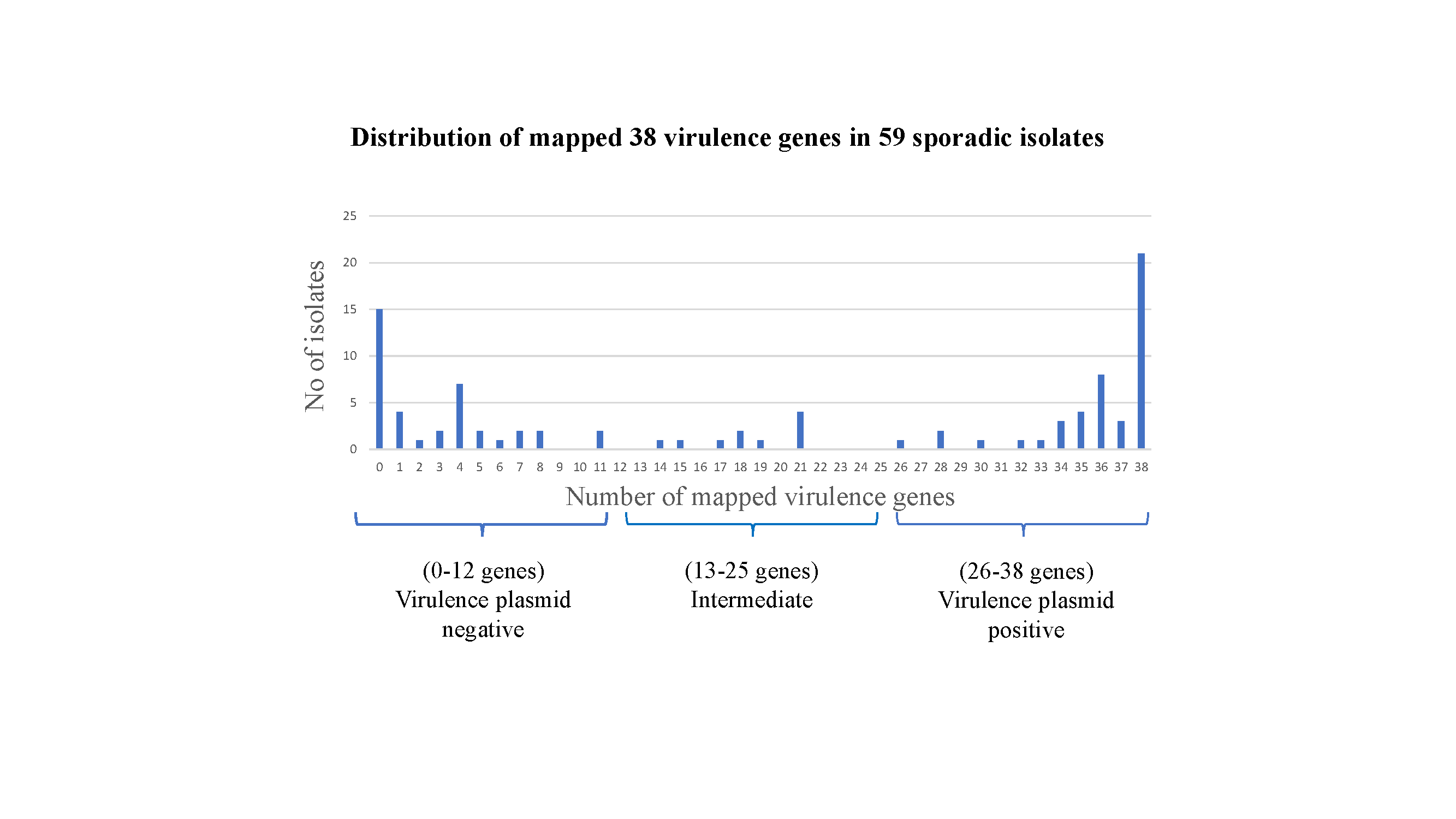

### Figure S4A.tiff

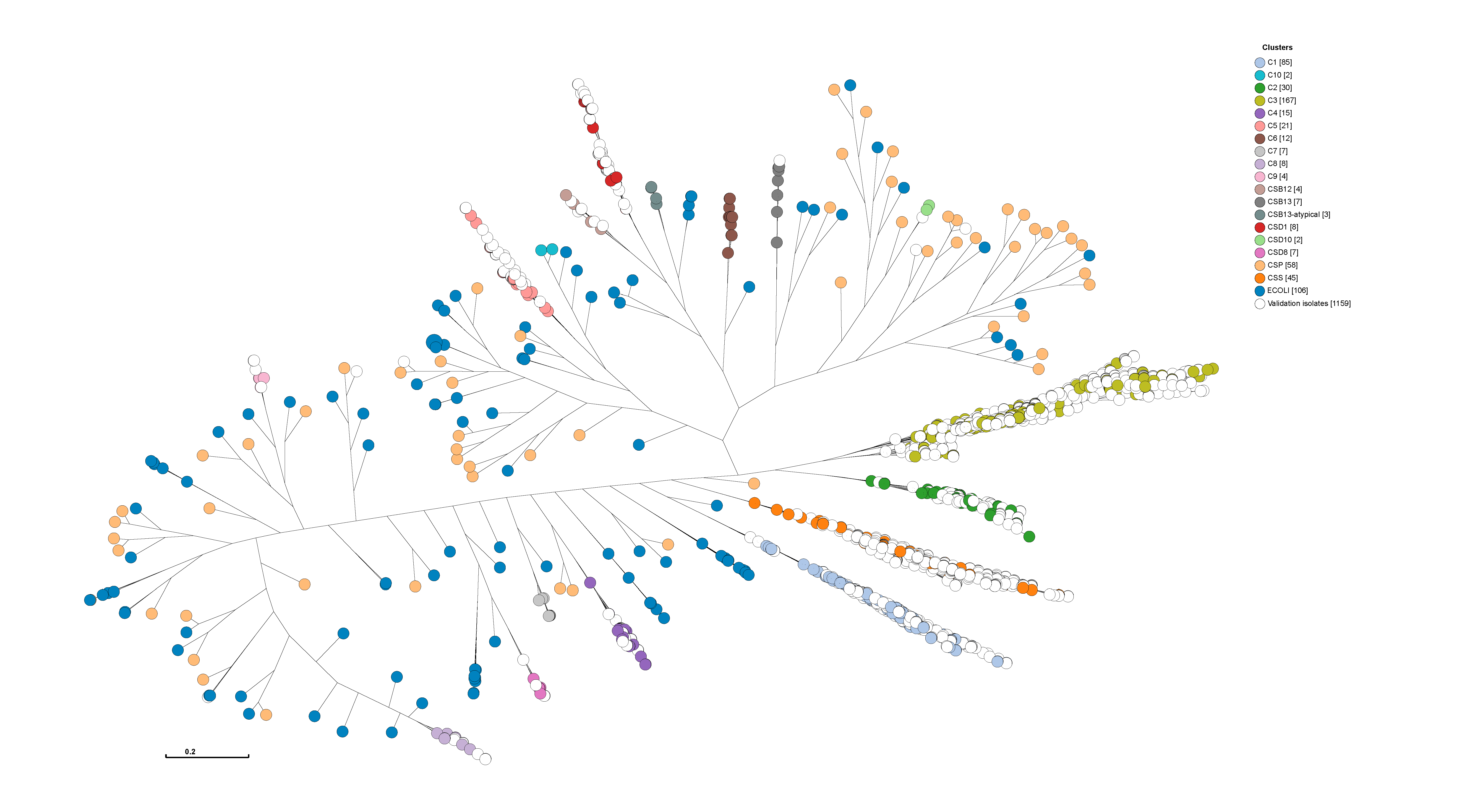

### Figure S4B.tiff

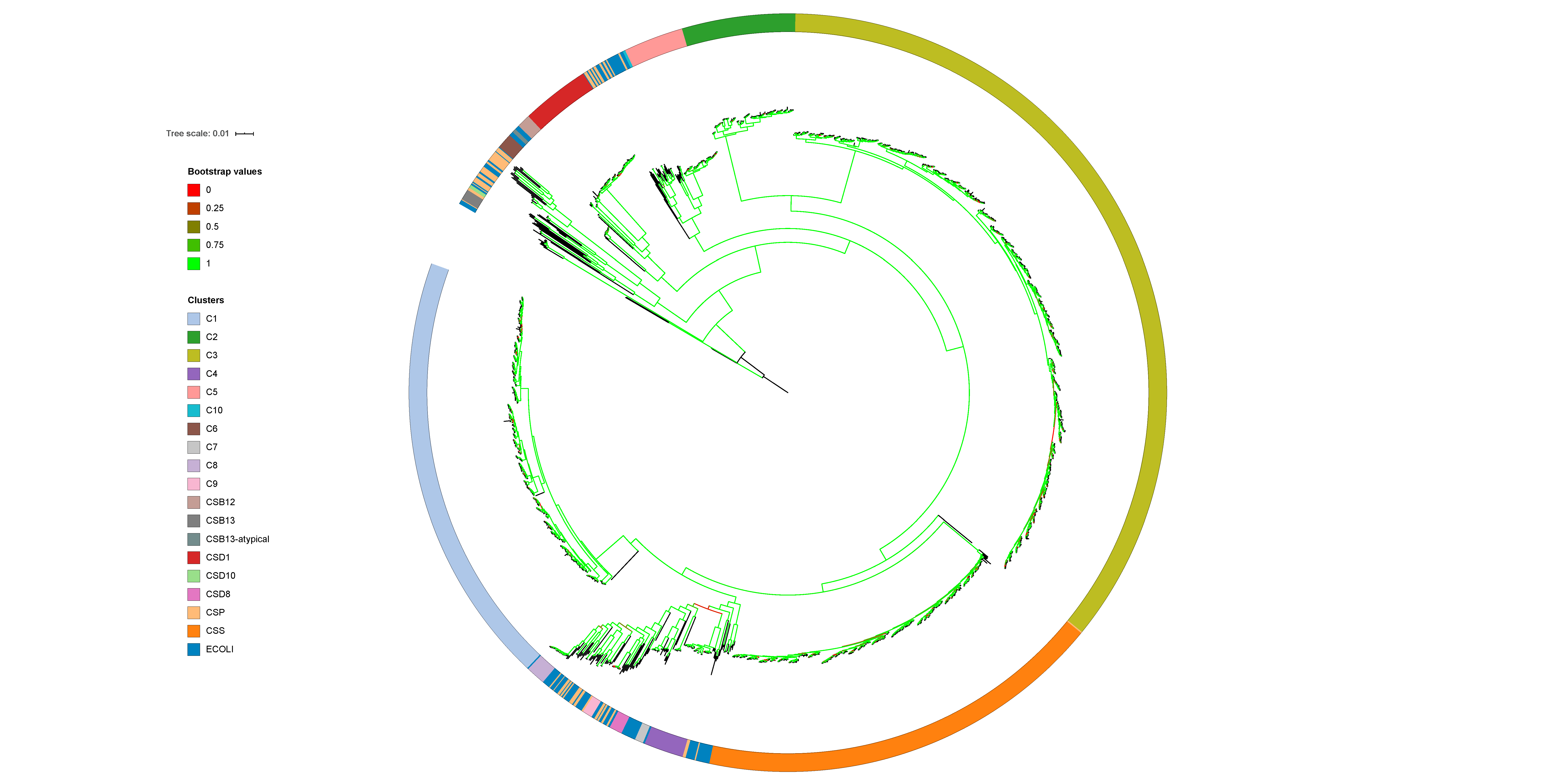
